# Convergent consequences of parthenogenesis on stick insect genomes

**DOI:** 10.1101/2020.11.20.391540

**Authors:** Kamil S. Jaron, Darren J. Parker, Yoann Anselmetti, Patrick Tran Van, Jens Bast, Zoé Dumas, Emeric Figuet, Clémentine M. François, Keith Hayward, Victor Rossier, Paul Simion, Marc Robinson-Rechavi, Nicolas Galtier, Tanja Schwander

**Affiliations:** Department of Ecology and Evolution, University of Lausanne, Lausanne, Switzerland; Swiss Institute of Bioinformatics, Lausanne, Switzerland; Institute of Evolutionary Biology, School of Biological Sciences, University of Edinburgh, Edinburgh, EH9 3FL; ISEM - Institut des Sciences de l’Evolution, Montpellier, France; CoBIUS lab, Department of Computer Science, University of Sherbrooke, Sherbrooke, Canada; Institute for Zoology, University of Cologne, Köln, Germany; Université Claude Bernard Lyon 1, CNRS, ENTPE, UMR 5023 LEHNA, F-69622, Villeurbanne, France; Université de Namur, LEGE, URBE, Namur, 5000, Belgium

**Author notes:** shared first authorship (Kamil S. Jaron, Darren J. Parker, Yoann Anselmetti). equal contribution (Marc Robinson-Rechavi, Nicolas Galtier, Tanja Schwander).

## Abstract

The shift from sexual reproduction to parthenogenesis has occurred repeatedly in animals, but how the loss of sex affects genome evolution remains poorly understood. We generated *de novo* reference genomes for five independently evolved parthenogenetic species in the stick insect genus *Timema* and their closest sexual relatives. Using these references in combination with population genomic data, we show that parthenogenesis results in an extreme reduction of heterozygosity, and often leads to genetically uniform populations. We also find evidence for less effective positive selection in parthenogenetic species, supporting the view that sex is ubiquitous in natural populations because it facilitates fast rates of adaptation. Contrary to studies of non-recombining genome portions in sexual species, genomes of parthenogenetic species do not accumulate transposable elements (TEs), likely because successful parthenogens derive from sexual ancestors with inactive TEs. Because we are able to conduct replicated comparisons across five species pairs, our study reveals, for the first time, how animal genomes evolve in the absence of sex in natural populations, providing empirical support for the negative consequences of parthenogenetic reproduction as predicted by theory.

## Introduction

Sex: What is it good for? The reason why most eukaryotes take a complicated detour to reproduction, when more straightforward options are available, remains a central and largely unanswered question in evolutionary biology (*1*, *2*). Animal species in which parthenogenetic reproduction is the sole form of replication typically occur at the tips of phylogenies and only a few of them have succeeded as well as their sexually reproducing relatives (*3*). In other words, most parthenogenetic lineages may eventually be destined for extinction. These incipient evolutionary failures, however, are invaluable as by understanding their fate something may be learned about the adaptive value of sex.

Parthenogenesis is thought to be favored in the short term because it generates a transmission advantage (*4*, *5*), as well as the advantage of assured reproduction when mates are scarce (*6*, *7*). The short-term benefits of parthenogenesis, however, are believed to come along with long-term costs. For example, the physical linkage between loci it entails can generate interferences that decrease the efficacy of natural selection (e.g. (*8*–*10*), reviewed in (*11*)). This is expected to translate into reduced rates of adaptation and increased accumulation of mildly deleterious mutations, which may potentially drive the extinction of parthenogenetic lineages.

In addition to these predicted effects on adaptation and mutation accumulation, parthenogenesis is expected to drive major aspects of genome evolution. A classical prediction is that heterozygosity (i.e., intra-individual polymorphism) increases over time in the absence of recombination, as the two haploid genomes diverge independently of each other, generating the so-called “Meselson Effect” (*12*, *13*). Parthenogenesis can also affect the dynamics of transposable elements (TEs), resulting in either increased or decreased genomic TE loads (*14*–*16*). Finally, some forms of parthenogenesis might facilitate the generation and maintenance of structural variants, which in sexuals are counter-selected due to the constraints of properly pairing homologous chromosomes during meiosis (*17*).

We tested these predictions by comparing the genomes of five independently derived parthenogenetic stick insect species in the genus *Timema* with their close sexual relatives (Figure 1). These replicate comparisons allowed us to solve the key problem in understanding the consequences of parthenogenesis for genome evolution: separating the consequences of parthenogenesis from lineage specific effects (*17*). *Timema* are wingless, plant-feeding insects endemic to western North America. Parthenogenetic species in this genus are diploid and of non-hybrid origin (*18*) and ecologically similar to their sexual relatives. Previous research, based on a small number of microsatellite markers, has suggested that oogenesis in parthenogenetic *Timema* is functionally mitotic, as no loss of heterozygosity between females and their offspring was detected (*18*).

**Figure 1.**
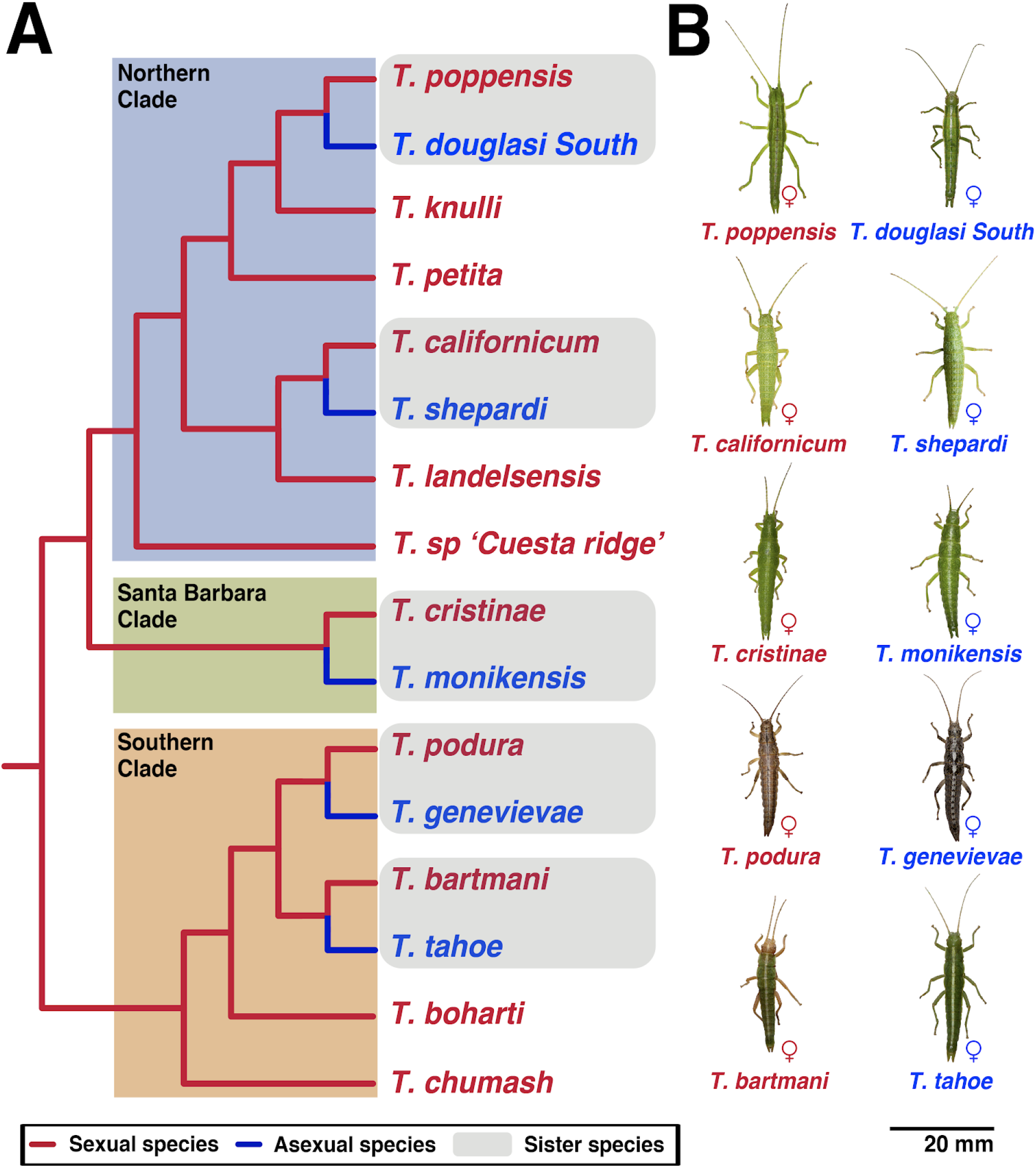
Multiple, independent transitions from sexual to parthenogenetic reproduction are known in the genus *Timema (19)*, each representing a biological replicate of parthenogenesis, and with a close sexual relative at hand for comparison **A.** Phylogenetic relationships of *Timema* species (adapted from (*19*, *20*)). **B.** Species sequenced in this study. Photos taken by © Bart Zijlstra - www.bartzijlstra.com.

### *De novo* genomes reveal extremely low heterozygosity in parthenogenetic stick insects

We generated ten *de novo* genomes of *Timema* stick insects, from five parthenogenetic and five sexual species (Figure 1, SM Tables 1, 2). Genomes were subjected to quality control, screened for contamination, and annotated (see Methods, SM text 1). The final reference genomes were largely haploid, spanned 75-95% of the estimated genome size (1.38 Gbp (*21*)), and were sufficiently complete for downstream analyses, as shown by the count of single copy orthologs conserved across insects (96% of BUSCO genes (*22*) detected on average; SM Table 3). A phylogeny based on a conservative set of 3975 1:1 orthologous genes (SM Table 4) corroborated published phylogenies and molecular divergence estimates in the *Timema* genus (SM Figure 1). Finally, we identified 55 putative horizontal gene acquisitions from non-metazoans, and they all happened well before the evolution of parthenogenesis (SM text 2).

We estimated genome-wide nucleotide heterozygosity in each reference genome directly from sequencing reads, using a reference-free technique (genome profiling analysis (*23*)). These analyses revealed extreme heterozygosity differences between the sexual and parthenogenetic species. The five sexual *Timema* featured nucleotide heterozygosities within the range previously observed in other sexual species (Figure 2; (*24*, *25*)). The heterozygosities in the parthenogenetic species were substantially lower, and in fact so low that reference-free analyses could not distinguish heterozygosity from sequencing error (SM text 3). We therefore compared heterozygosity between sexuals and parthenogens by calling SNPs in five re-sequenced individuals per species. This analysis corroborated the finding that parthenogens have extremely low (<10^−5^) heterozygosity, being at least 140 times lower than that found in their sexual sister species (permutation ANOVA, reproductive mode effect p = 0.0049; Figure 2). Screening for structural variants (indels, tandem duplications, and inversions) in sexual and parthenogenetic individuals revealed the same pattern: extensive and variable heterozygosity in sexual species and homozygosity in the parthenogens (Figure 2, SM text 3). Some heterozygosity in *Timema* parthenogens could be present in genomic regions not represented in our assemblies, such as centromeric and telomeric regions. These regions however represent a relatively small fraction of the total genome, meaning that for most of the genome at least, *Timema* parthenogens are either largely or completely homozygous for all types of variants (SM text 3).

**Figure 2.**
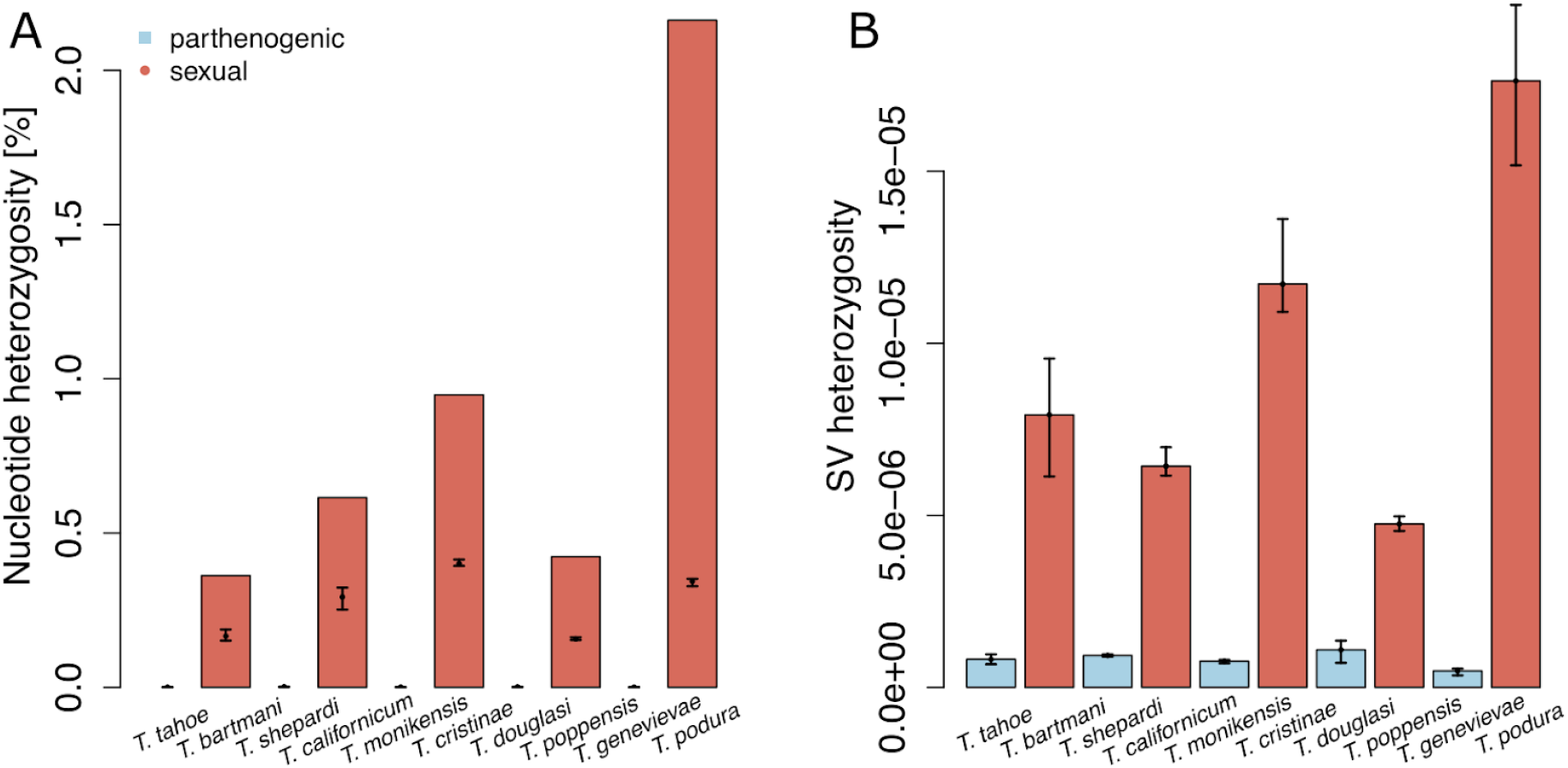
Extremely low heterozygosity in parthenogenetic *Timema* species for different types of variants. **A.** Nucleotide heterozygosity represented by bars indicates genome-wide estimates for the reference genomes (based on raw reads, see Methods), heterozygosity based on SNP calls in re-sequenced individuals is indicated by points and represents a conservative estimation of heterozygosity in the assembled genome portions (with error bars indicating the range of estimates across individuals) **B.** Heterozygous structural variants (SVs, reported as number of heterozygous SVs / number of callable sites) in re-sequenced individuals (with error bars indicating the range of estimates across individuals). Note that even though heterozygous SNPs and SVs were called using stringent parameters, it is likely that a large portion are false positives in parthenogenetic *Timema* (see SM text 3).

The unexpected finding of extremely low heterozygosity in *Timema* parthenogens raises the question of when and how heterozygosity was lost. For example, the bulk of heterozygosity could have been lost during the transition from sexual reproduction to parthenogenesis (*26*). Alternatively, heterozygosity loss could be a continuous and ongoing process in the parthenogenetic lineages. To distinguish these options, we investigated the origin of the genetic variation present among different homozygous genotypes in each parthenogenetic species. We found that only 6-19% of the SNPs called in a parthenogen are at positions that are also polymorphic in the sexual relative (SM Table 5). This means that most of the variation in parthenogens likely results from mutations that appeared after the split from the sexual lineage. This implies that heterozygosity generated through new mutations is lost continuously in parthenogens, and was not suddenly lost at the inception of parthenogenesis. The most likely explanation for these findings is that parthenogenetic *Timema* are, in fact, not functionally mitotic but automictic. Automictic parthenogenesis frequently involves recombination and segregation, and can lead to homozygosity in most or all of the genome (*27*, *28*). Although automixis can allow for the purging of heterozygous deleterious mutations (*29*), the classical predictions for the long-term costs of asexuality extend to automictic parthenogens because, as for obligate selfers, linkage among genes is still much stronger than in classical sexual species (*30*). This is especially the case in largely homozygous parthenogens, where recombination and segregation, even if mechanistically present, have no effect on genotype diversities.

Functional mitosis in *Timema* was previously inferred from the inheritance of heterozygous microsatellite genotypes between females and their offspring (*18*), a technique widely used in non-model organisms with no cytological data available (e.g., (*31*, *32*)). The most likely reconciliation of these contrasting results is that heterozygosity is maintained in only a small portion of the genome, for example the centromeres or telomeres, or between paralogs. Consistent with this idea, we were unable to locate several of the microsatellite-containing regions in even the best *Timema* genome assemblies (SM text 4), suggesting that these regions are not present in our assemblies due to the inherent difficulty of assembling repetitive genome regions from short read data (*33*).

### Extensive variation in genotype diversity between parthenogenetic populations

Parthenogenesis and sexual reproduction are expected to drive strikingly different distributions of polymorphisms in genomes and populations. Different regions within genomes experience different types of selection with sometimes opposite effects on the levels of polymorphisms within populations, such as purifying versus balancing selection (*34*). The increased linkage among genes in parthenogenetic as compared to sexual species is expected to homogenize diversity levels across different genome regions. Furthermore, recurrent sweeps of specific genotypes in parthenogenetic populations can lead to extremely low genetic diversity and even to the fixation of a single genotype, while sweeps in sexual populations typically reduce diversity only in specific genome regions.

To address these aspects in the genomes of sexual and parthenogenetic *Timema* species, we mapped population-level variation for the SNPs and SVs inferred above to our species-specific reference genomes. We then anchored our reference genome scaffolds to the 12 autosomal linkage groups of a previously published assembly of the sexual species *T. cristinae* (v1.3 from (*35*), SM text 5). This revealed that different types of polymorphisms (SNPs and SVs) tended to co-occur across the genomes in all species, independently of reproductive mode (Figure 3).

**Figure 3.**
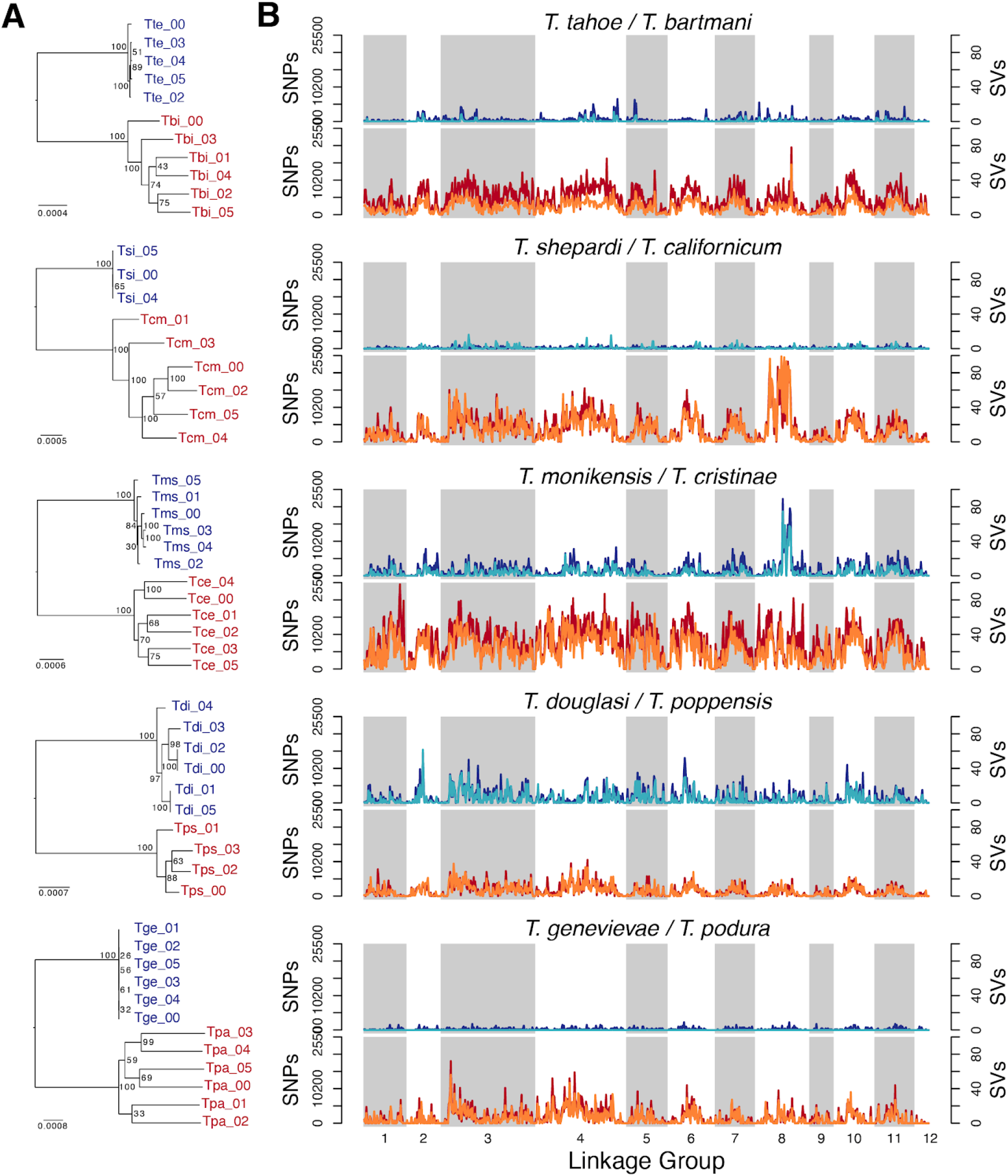
Population polymorphism levels in parthenogenetic (blue) and sexual (red) *Timema* species. **A.** Phylogenies based on 1:1 orthologous genes reflect the different levels of genotype diversities in parthenogenetic *Timema* species **B.** Distribution of structural variants (SVs; dark blue and red) and SNPs (light blue and orange) along the genome. Scaffolds from the ten *de novo* genomes are anchored on autosomal linkage groups from the sexual species *T. cristinae* (SM text 5).

The focal population for three of the five parthenogenetic species (*T. genevievae*, *T. tahoe* and *T. shepardi*) consisted largely of a single genotype with only minor variation among individuals. By contrast, genotype diversity was considerable in *T. monikensis* and *T. douglasi* (Figure 3A). In the former species, there was further a conspicuous diversity peak on LG8, supporting the idea that parthenogenesis is automictic in *Timema*. Indeed, under complete linkage (functionally mitotic parthenogenesis), putative effects of selection on this LG would be expected to propagate to the whole genome. Independently of local diversity peaks, overall diversity levels in *T. monikensis* and *T. douglasi* were comparable to the diversities in populations of some of the sexual *Timema* species (Figure 3A). Different mechanisms could contribute to such unexpected diversities in parthenogenetic *Timema*, including the presence of lineages that derived independently from their sexual ancestor, or rare sex. While a single transition to parthenogenesis is believed to have occurred in *T. monikensis*, the nominal species *T. douglasi* is polyphyletic and known to consist of independently derived clonal lineages. These lineages have broadly different geographic distributions but can overlap locally (*19*). Identifying the causes of genotypic variation in these species, including the possibility of rare sex, requires further investigation and is a challenge for future studies.

Independently of the mechanisms underlying polymorphism in the parthenogenetic species *T. monikensis*, the polymorphism peak on LG8 is striking (Figure 3B). This peak occurs in a region previously shown to determine color morph (green, green-striped, or brownish (“melanistic”)) in the sexual sister species of *T. monikensis*, *T. cristinae (35)*. Our focal *T. monikensis* population features four discrete color morphs (green, dark brown, yellow, and beige), suggesting that additional color morphs may be regulated by the region identified in *T. cristinae*. We also found a peak in polymorphism on LG8, spanning over approximately two-thirds of LG8, in the sexual species *T. californicum*, which features a different panel of color morphs than *T. cristinae (36)*. Interestingly, this diversity peak in *T. californicum* was generated by the presence of two divergent haplotypes (approximately 24Mbp long), with grey individuals homozygous for one haplotype and green individuals heterozygous or homozygous for the alternative haplotype (SM text 6). Note that the grey color morph is not known in the monomorphic green parthenogenetic sister of *T. californicum* (*T. shepardi*), and we therefore do not expect the same pattern of polymorphism on LG8 in this species.

### Faster rate of adaptive evolution in sexual than parthenogenetic species

We have shown previously that parthenogenetic *Timema* species accumulate deleterious mutations faster than sexual species (*37*, *38*), a pattern also reported in other parthenogenetic taxa (reviewed in (*17*, *39*)). This is expected given that linkage among loci in parthenogens prevents selection from acting individually on each locus, which generates different forms of selective interference (*9*, *10*, *40*). In addition to facilitating the accumulation of deleterious mutations, selective interference among loci in parthenogens should also constrain the efficiency of positive selection. While there is accumulating evidence for this process in experimental evolution studies (e.g., (*41*–*43*)), its impact on natural populations remains unclear (*17*, *39*). To compare the efficiency of positive selection in sexual and parthenogenetic *Timema*, we used a branch-site model on the gene trees ((*44*), Methods). We compared the terminal branches leading to sexual or parthenogenetic species in one-to-one orthologous genes identified in at least three species pairs (SM Table 4), using a threshold of q < 0.05 to classify which terminal branches show evidence of positive selection.

We found a greater number of positively selected genes in sexual than parthenogenetic species (Figure 4, binomial GLMM p = 0.005). In addition, we also examined if there was more evidence for positive selection in sexual species in a threshold-free way by comparing the likelihood ratio test statistic between parthenogenetic and sexual species (as in (*45*, *46*)). This confirmed that the evidence for positive selection was stronger for sexual species (permutation glm p = 0.011).

**Figure 4.**
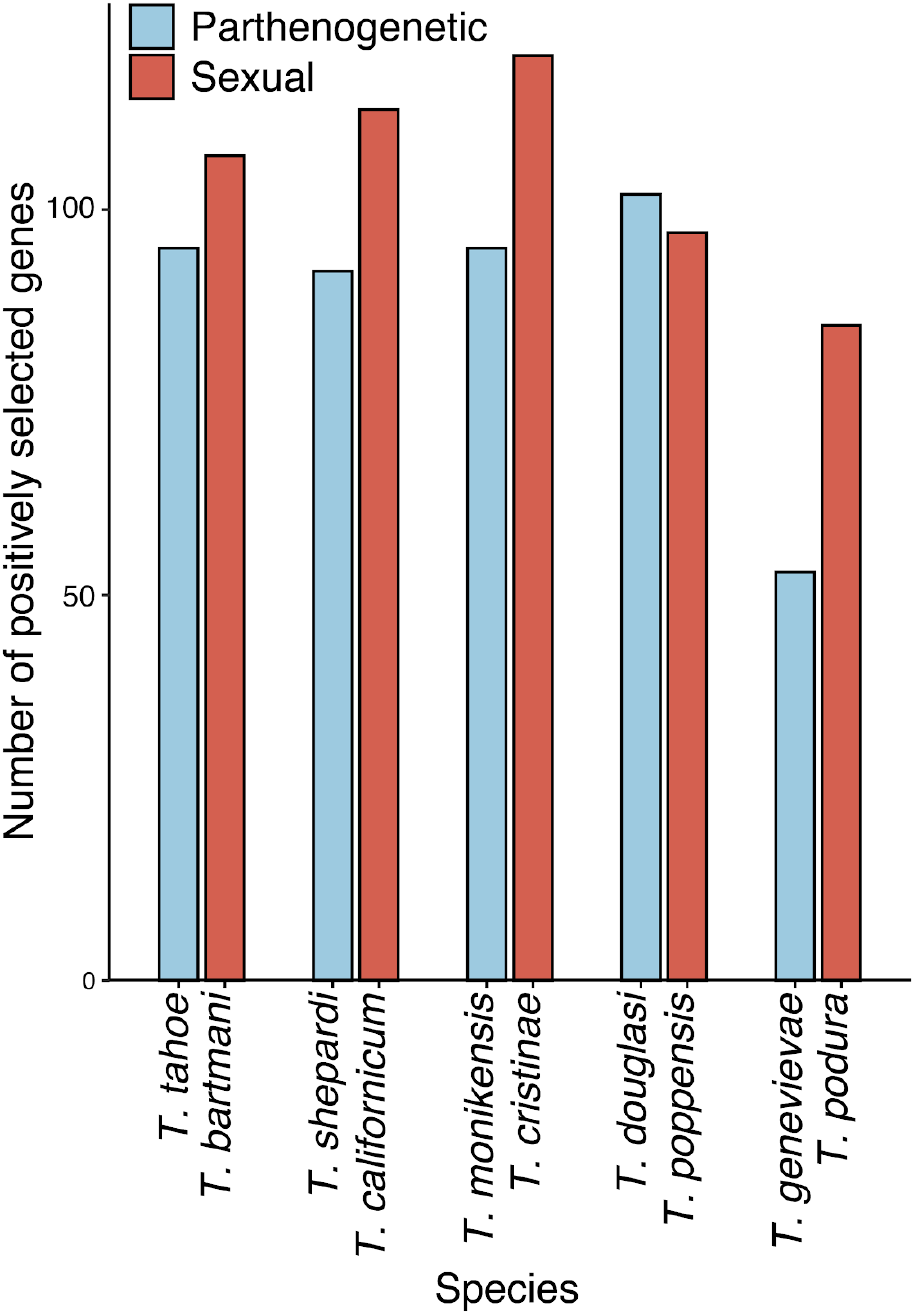
Number of genes showing evidence for positive selection in each species. In addition to reproductive mode, species pair also had a significant influence on the number of positively selected branches (binomial GLMM p = 0.015). There was no significant interaction between species pair and reproductive mode (p = 0.197). Note, the difference between reproductive modes is robust to a more stringent cutoff (q < 0.01 instead of 0.05, SM Figure 2).

The positively selected genes we identified are most likely associated with species-specific adaptations. Few of them were shared between species, with overlap between species not greater than expected by chance (SM Figure 3, FDR < 0.4), and there was little enrichment of functional processes in positively selected genes (0-19 GO terms per species, SM Table 8). Interestingly, most of the significant GO terms were associated with positively selected genes in parthenogenetic *Timema* (SM Table 8), likely because a much smaller proportion of positively selected genes in sexual species had annotations (SM Figure 4). We speculate that positively selected genes in sexuals could often be involved in sexual selection and species recognition. Indeed, genes associated with processes such as pheromone production and reception often evolve very fast, which makes them difficult to annotate through homology-based inference (*47*). For the parthenogenetic species, although some terms could be associated with their mode of reproduction (e.g. GO:0033206 meiotic cytokinesis in *T. douglasi*), most are not clearly linked to a parthenogenetic life cycle.

### Transposable element loads are similar between species with sexual and parthenogenetic reproduction

Upon the loss of sexual reproduction, transposable element (TE) dynamics are expected to change (*14*, *16*, *48*). How these changes affect genome-wide TE loads is however unclear as sex can facilitate both the spread and the elimination of TEs (*17*). In parthenogens, TE load might initially increase as a result of weaker purifying selection, a pattern well illustrated by the accumulation of TEs in non-recombining parts of sex chromosomes and other supergenes (*49*, *50*). However, TE loads in parthenogens are expected to decrease over time via at least two non-mutually exclusive mechanisms. First, TEs are expected to evolve lower activity over time as their evolutionary interests are aligned with their hosts (*14*, *48*). Second, TE copies that were purged via excision can re-colonize a sexual but not a parthenogenetic genomic background (*15*, *16*). Finally, it is important to note that the predicted effects of reproductive mode on TE loads require some amount of TE activity (active transposition or excision) to occur. Without such activity, TE content does not vary among individuals and can therefore not change over time.

We generated a *Timema* genus-level TE library by merging *de novo* TE libraries generated separately for each of the ten *Timema* species. We then quantified TE loads in each *Timema* genome by mapping reads to this merged library (see **Methods**). The overall TE content was very similar in all ten species (20 - 23.6%), with significant differences in abundance of TE superfamilies between species groups but no significant effect of reproductive mode (p=0.43; Figure 5; SM Figure 5).

**Figure 5.**
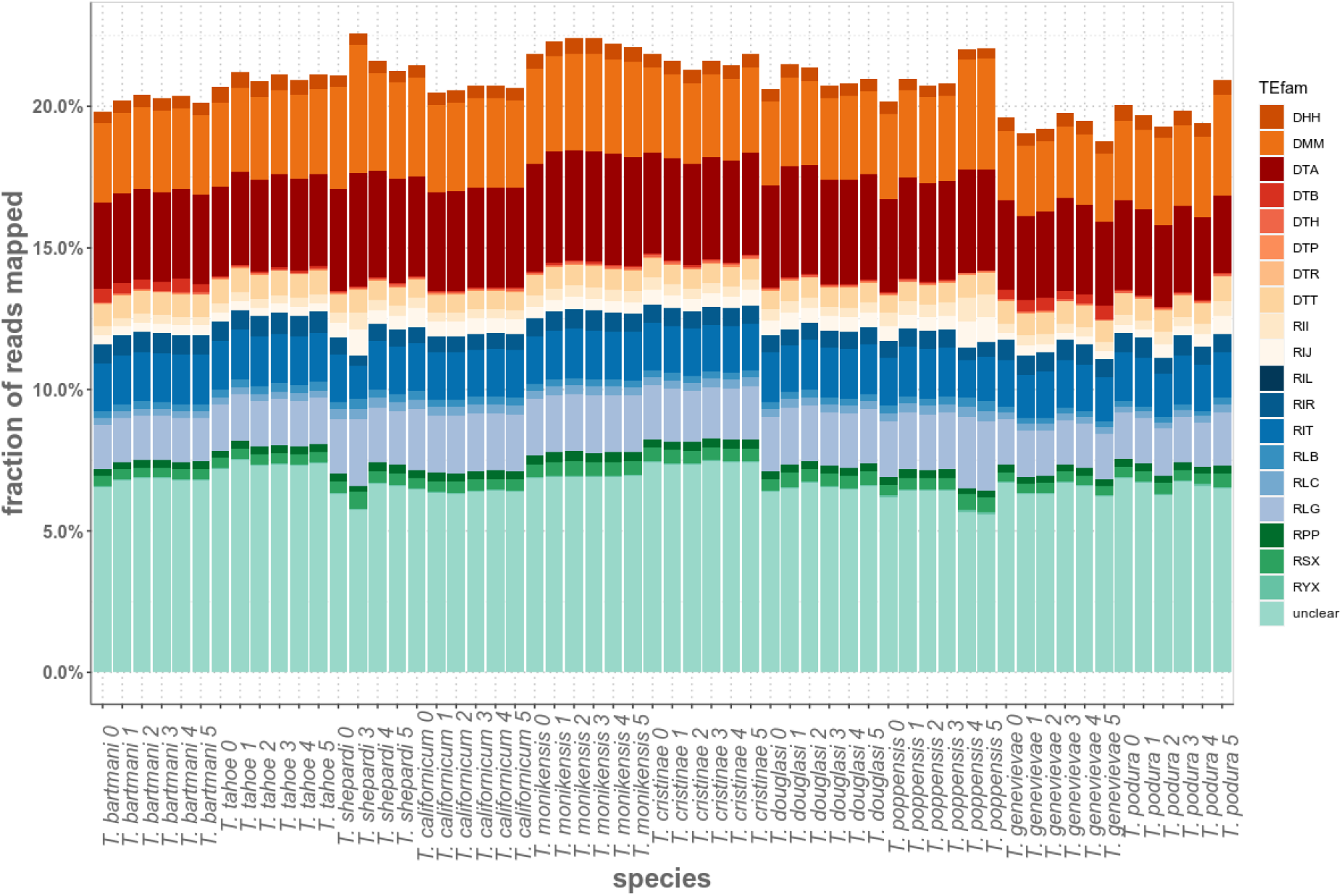
Total TE abundance in the ten *Timema* species. TE abundance is expressed as the fraction of reads that map to a genus-level TE library. TE families are named following the Wicker classification (*51*). The first character corresponds to the TE class (Class I are retrotransposons (R), Class II are DNA transposons (D)), the second character corresponds to the Order (e.g. LTR) and the third to the Superfamily (e.g. *Gypsy*); for example, RLG is a *Gypsy* retroelement. The character X indicates unknown classification at the superfamily level (because of fragmentation or lack of detectable homology).

No difference in TE load between sexual and parthenogenetic *Timema* would be expected if TEs were already well controlled in their ancestor, without any subsequent TE activity. Consistent with this idea, we find very little evidence for ongoing TE activity in the genus. The oldest node in our *Timema* phylogeny has an age estimate of 30 Mya (*20*) but the TE contents of the two clades separating at this node have only diverged by 1.3%, suggesting that TEs remained largely silent during the evolution of the genus. Inactive TEs may facilitate the persistence of incipient parthenogenetic strains (*17*) and thus help to explain the high frequency of established parthenogenetic species in *Timema*.

## Conclusion

We present genomes of five independently derived parthenogenetic lineages of *Timema* stick insects, together with their five sexual sister species. This design with replicated species pairs allows us, for the first time, to disentangle consequences of parthenogenesis from species-specific effects. All parthenogenetic *Timema* species are largely or completely homozygous for both SNPs and SVs, and frequently feature lower levels of population polymorphism than their close sexual relatives. Low population polymorphism can exacerbate the effects of linkage for reducing the efficacy of selection, resulting in reduced rates of positive selection in parthenogenetic *Timema*, in addition to the accumulation of deleterious mutations previously documented (*37*). In spite of these negative genomic consequences, parthenogenesis is an unusually successful strategy in *Timema*. It evolved and persisted repeatedly in the genus, and parthenogenetic species often occur across large geographic areas. Because *Timema* are wingless and their populations subjected to frequent extinction-recolonization dynamics in their fire-prone Californian shrubland habitats, the genomic costs of parthenogenesis are likely offset by one of the most classical benefits of parthenogenesis: the ability to reproduce without a mate.

## Methods

### Sample collection and sequencing

For each of the ten species, the DNA for Illumina shotgun sequencing was derived from virgin adult females collected in 2015 from natural populations in California (SM Table 1). Extractions were done using the Qiagen Mag Attract de HMW DNA kit, following manufacturer indications. Five PCR-free libraries were generated for each reference genome (three 2×125bp paired end libraries with average insert sizes of respectively 350, 550 and 700bp, and two mate-pair libraries with 3000 and 5000bp insert sizes), one library (550bp insert size) was generated for each re-sequenced individual. Libraries were prepared using the illumina TruSeq DNA PCR-Free or Nextera Mate Pair Library Prep Kits, following manufacturer instructions, and sequenced on the Illumina HiSeq 2500 system, using v4 chemistry and 2× 125 bp reads at FASTERIS SA, Plan-les-Ouates, Switzerland.

### Genome assembly and annotation

The total coverage for the reference genomes (all libraries combined) ranged between 37-45× (SM Table 2). Trimmed paired-end reads were assembled into contigs using ABySS (*52*) and further scaffolded using paired-end and mate pairs using BESST (*53*). Scaffolds identified as contaminants were filtered using Blobtools (*54*). The assembly details can be found in supplementary materials (SM text 1).

Publically available RNA-seq libraries for *Timema (37, 55, 56)* were used as expression evidence for annotation. Trimmed reads were assembled using Trinity v2.5.1 (*57*) to produce reference-guided transcriptomes. The transcriptomes and protein evidence were combined with *ab initio* gene finders to predict protein coding genes using MAKER v2.31.8 (*58*). The annotation details can be found in the supplementary materials (SM text 1).

### Orthologs

*Timema* orthologous groups (OGs) were inferred with the OrthoDB standalone pipeline (v. 2.4.4) using default parameters (*59*). In short, genes are clustered with a graph-based approach based on all best reciprocal hits between each pair of genomes. The high level of fragmentation typical for Illumina-based genomes constrains the ability to identify 1:1 orthologs across all ten *Timema* species. To maximize the number of single copy OGs covering all ten *Timema* species, transcriptomes were included during orthology inference. Thus, transcripts were used to complete OGs in absence of a gene from the corresponding species. Using this approach, 7157 single copy OGs covering at least three sexual-parthenogenetic sister species pairs were obtained (SM Table 4).

### Horizontal gene transfers (HGT)

To detect HGT from non-metazoan species, we first used the pipeline of foreign sequence detection developed by Francois et al. (*60*). We used the set of CDS identified in publicly available transcriptomes (*37*) and the genome assemblies prior to the decontamination procedure with Blobtools (*54*). The rationale is that some genuine HGT could have been wrongly considered as contaminant sequences during this decontamination step and thus been removed from the assembly. Scaffolds filtered during decontamination are available from our github repository (https://github.com/AsexGenomeEvol/Timema_asex_genomes/tree/main/4_Horizontal_Gene_Transfers/contamination_sequences), and will be archived upon acceptance.

Briefly, a DIAMOND BlastP (v0.8.33) (*61*) allows to detect candidate non-metazoan genes in the set of CDS of each species. Taxonomic assignment is based on the 10 best blast hits to account for potential contaminations and other sources of taxonomic misassignment in the reference database. Candidate non-metazoan sequences are then subjected to a synteny-based screen with Gmap (v2016-11-07) (*62*) to discriminate between contaminant sequences and potential HGT-derived sequences. A sequence is considered as a HGT candidate if it is physically linked to (i.e., mapped to the same scaffold as) at least one “confident-arthropod” CDS (previously identified in the DIAMOND blast).

We then clustered all HGT candidates identified in each of the 10 *Timema* species into HGT families using Silix (v1.2.10) (*63*), requiring a minimum of 85% identity (default parameters otherwise). These HGT families were then “completed” as much as possible by adding homologs from the genome assemblies not identified as HGT candidates (this could occur if the corresponding sequences are fragmented or on short scaffolds for example). To this end, the longest sequence of each HGT family was mapped (using Gmap) on the genomic scaffolds of all species, requiring a minimum of 85% identity.

For each completed HGT family, a protein alignment of the candidate HGT sequence(s) and its (their) 50 best DIAMOND blastP hits in the reference database (1^st^ step of the pipeline) was generated with MAFFT (v7) (*64*). The alignments were cleaned using HMMcleaner (stringency parameter = 12) (*65*) and sites with more than 50% missing data were removed. Phylogenetic trees were inferred using RAxML (v8.2) (*66*) with the model ‘PROTGAMMALGX’ of amino-acid substitution and 100 bootstrap replicates. Phylogenetic trees were inspected by eye to confirm or not an evolutionary history consistent with the hypothesis of HGT.

### Heterozygosity

Genome-wide nucleotide heterozygosity was estimated using genome profiling analysis of raw reads from the reference genomes using GenomeScope (v2) (*23*). A second, SNP-based heterozygosity estimate was generated using re-sequenced individuals. We re-sequenced five individuals per species, but 3 individuals of *T. shepardi*, 2 individuals of *T. poppensis* and one *T. tahoe* individual did not pass quality control and were discarded from all downstream analyses. SNP calling was based on the GATK best practices pipeline (*67*). We used a conservative set of SNPs with quality scores ≥300, and supported by 15x coverage in at least one of the individuals. SNP heterozygosity was then estimated as the number of heterozygous SNPs divided by the number of callable sites in each genome. Due to stringent filtering criteria, our SNP based heterozygosity is an underestimation of genome-wide heterozygosity.

### Structural variants

We used Manta (v1.5.0) (*68*), a diploid-aware pipeline for structural variant (SV) calling, in the same set of re-sequenced individuals used for SNP heterozygosity estimates. We found a high frequency of heterozygous SVs with approximately twice the expected coverage (SM Figure 7), which likely represent false positives. To reduce the number of false positives, we filtered very short SVs (30 bases or less) and kept only variant calls that had either split read or paired-end read support within the expected coverage range, where the coverage range was defined individually for each sample by manual inspection of coverage distributions. The filtered SV calls were subsequently merged into population SV calls using SURVIVOR (v1.0.2) (*69*). The merging criteria were: SV calls of the same type on the same strand with breakpoints distances shorter than 100 bp.

### Genome alignment

We anchored our genome assemblies to the reference of *T. cristinae* (BioProject Accession PRJNA417530) (*35*) using MUMmer (version 4.0.0beta2) (*70*) with parameter --mum. The alignments were processed by other tools within the package: show-coords with parameters -THrcl to generate tab-delimited alignment files and dnadiff to generate 1-to-1 alignments. We used only uniquely anchored scaffolds for which we were able to map at least 10k nucleotides to the *T. cristinae* reference genome.

### Transposable elements

For each species, specific repeat libraries were constructed and annotated to the TE superfamily level (*51*) wherever possible. For collecting repetitive sequences, we used a raw read based approach DNAPipeTE v1.2 (*71*) with parameters-genome_coverage 0.5-sample_number 4 and respective species genome size, as well as an assembly based approach (RepeatModeler v1.0.8 available at http://www.repeatmasker.org/RepeatModeler/), such that repeats not present in the assembly can still be represented in the repeat library. The two raw libraries were merged and clustered by 95% identity (the TE family threshold) using usearch v10.0.240 (*72*) with the centroid option. To annotate TEs larger than 500 bp in the repeat library, we used an approach that combines homology and structural evidence (PASTEClassifier (*73*)). Because PASTEClassifier did not annotate to TE superfamily levels, we additionally compared by BlastN (v. 2.7.1+) (*74*) the repeat libraries to the well curated *T. cristinae* TE library from Soria-Carrasco et al. (*21*). Blast hits were filtered according to TE classification standards: identity percentage >80%, alignment length >80 bp, and the best hit per contig was kept. The two classification outputs were compared and in case of conflict the classification level of PASTEClassifier was preferred. All non-annotated repeats were labelled ‘unknown’. Repeat library header naming was done according to RepeatMasker standard, but keeping the Wicker naming for elements (i.e., Wicker#Repeatmasker, e.g., DTA#DNA/hAT). TE libraries were sorted by header and TE annotations to similar families numbered consecutively. Species-specific TE libraries were merged into a genus-level *Timema* TE library to account for any TE families that might have not been detected in the single species assemblies.

To estimate the TE load of reference genomes and resequenced individuals, we first repeat masked the assemblies with the genus-level TE library using RepeatMasker v4.1.0 with parameters set as -gccalc -gff -u -a -xsmall -no_is -div 30 -engine rmblast (*75*). Second, we mapped the 350 bp insert paired-end reads back to the reference genome assemblies using BWA-MEM v0.7.17 (*76*) with standard parameters. We then counted the fraction of reads mapping to TEs out of total mappable reads by counting the number of reads that mapped to each genomic location annotated as TE using htseq-counts (v0.6.1.1p1) (*77*) with parameters set to -r name -s no -t similarity -i Target --nonunique using the mapped read alignments and the gff output of RepeatMasker (filtered for TE length of >80 bp). TE loads were compared among species using a permutation ANOVA with 5000 bootstrap replicates.

### Positive selection analysis

Only one-to-one orthologs in at least three pairs of species (sister-species sex-asex) were used. The species phylogeny was imposed on every gene as the “gene tree”. We used a customized version of the Selectome pipeline (*78*). All alignment building and filtering was performed on predicted amino acid sequences, and the final amino acid MSAs (multiple sequence alignments) were used to infer the nucleotide MSAs used for positive selection inference. MSAs were obtained by MAFFT (v. 7.310) (*64*) with the allowshift option, which avoids over-aligning non homologous regions (e.g. gene prediction errors, or alternative transcripts). All the next steps “mask” rather than remove sites, by replacing the amino acid with a ‘X’ and the corresponding codon with ‘NNN’. MCoffee (v11.00.8cbe486) (*79*) was run with the following aligners: mafft_msa, muscle_msa, clustalo_msa (*80*), and t_coffee_msa (*81*). MCoffee provides a consistency score per amino acid, indicating how robust the alignment is at that position for that sequence. Residues with a consistency score less than 5 were masked. TrimAl (v. 1.4.1) (*82*) was used to mask columns with less than 4 residues (neither gap nor ‘X’).

The branch-site model with rate variation at the DNA level (*44*) was run using the Godon software (https://bitbucket.org/Davydov/godon/, version 2020-02-17, option BSG --ncat 4). Each branch was tested iteratively, in one run per gene tree. For each branch, we obtain a ΔlnL which measures the evidence for positive selection, a corresponding p-value and associated q-value (estimated from the distribution of p-values over all branches of all genes), and an estimate of the proportion of sites under positive selection if any. All positive selection results, and detailed methods, will be available at https://selectome.org/timema. To determine if there the number of positively selected genes differed between sexual than parthenogenetic species we used a binomial GLMM approach (lme4 (*83*)) with q-value threshold of 0.05 or 0.01. Significance of model terms was determined with a Wald statistic. In addition, we also examined if there was more evidence for positive selection in sexual species in a threshold-free way by comparing ΔlnL values between parthenogenetic and sexual species (as in (*45*, *46*)). To do this we used a permutation glm approach where reproductive mode (sexual or parthenogenetic) was randomly switched within a species-pair. To determine if the overlap of positively selected genes was greater than expected by chance we used the SuperExactTest package (v. 0.99.4) (*84*) in R. The resulting p-values were multiple test corrected using Benjamini and Hochberg’s algorithm implemented in R. Functional enrichment analyses were performed using TopGO (v. 2.28.0) (*85*) using the *D. melanogaster* functional annotation (see SM text 1). To determine if a GO term was enriched we used a Fisher’s exact test with the ‘weight01’ algorithm to account for the GO topology. GO terms were considered to be significantly enriched when p < 0.05.

## Data and code availability

Raw sequence reads have been deposited in NCBI’s sequence read archive under the following bioprojects: PRJNA371785 (reference genomes, SM Table 7A), PRJNA670663 (resequenced individuals, SM Table 7B), and PRJNA673001 (PacBio reads for *T. douglasi*). Genome assemblies and annotations PRJEB31411. Scripts for the analyses in this paper are available at: https://github.com/AsexGenomeEvol/Timema_asex_genomes. Data were processed to generate plots and statistics using R v3.4.4.

## Acknowledgements

We thank Bart Zijlstra, Chloé Larose, and Kirsten Jalvingh for help in the field, and Deborah Charlesworth, Daniel Wegmann, and current and previous members of the Schwander and Robinson-Rechavi labs for discussions. This project was supported by several grants from the Swiss Science Foundation (CRSII3_160723 as well as PP00P3_170627, 31003A_182495, IZLRZ3_163872, and 407540_167276) and funding from the University of Lausanne.

## Notes

### Competing Interest Statement

The authors have declared no competing interest.

### Summary of Updates

Minor edits

