## Supplemental tables and figures for "Convergent consequences of parthenogenesis on stick insect genomes"

### Supplementary materials: Convergent consequences of parthenogenesis on stick insect genomes

#### SM Table 1. Origin of biological material

All six females per species were taken from a single location at the indicated coordinates. Red species reproduce sexually (s), blue species via parthenogenesis (p).

| Species | Host plant | Coordinates | |
|---|---|---|---|
| | | longitude | latitude |
| *T. tahoe* (p) | *Abies concolor* | 38.7610110 | -120.1600530 |
| *T. bartmani* (s) | *Abies concolor* | 34.1700000 | -117.0020167 |
| *T. shepardi* (p) | *Arctostaphylos sp.* | 39.1926500 | -123.2617833 |
| *T. californicum* (s) | *Quercus sp.* | 37.3431667 | -121.6364667 |
| *T. douglasi* (p) | *Pseudotsuga menziesii* | 38.9825500 | -123.4697500 |
| *T. poppensis* (s) | *Sequoia sempervirens* | 37.1655167 | -122.0155500 |
| *T. monikensis* (p) | *Cercocarpus betuloides* | 34.1148833 | -118.8531333 |
| *T. cristinae* (s) | *Cercocarpus betuloides* | 34.5362700 | -119.2444300 |
| *T. genevievae* (p) | *Adenostoma fasciculatum* | 38.9957833 | -122.9257667 |
| *T. podura* (s) | *Adenostoma fasciculatum* | 33.7976020 | -116.7769850 |

**SM Table 2. Sequencing coverage**

Read coverage for the reference assemblies of individual *Timema* species was estimated using the haploid genome size of *Timema cristinae* of 1.381Gbp (*21*). Red species are reproducing sexually, while blue species are parthenogenetic. Is: insert size [bp].

| species | paired-end | | | mate-pair | | orphans | Total |
|---|---|---|---|---|---|---|---|
| | Is 350 | Is 550 | Is 700 | Is 3000 | Is 5000 | | |
| *T. tahoe* | 15 | 12.2 | 5.7 | 4.1 | 3 | 3 | 43.1 |
| *T. bartmani* | 12.3 | 13.5 | 3.7 | 2.7 | 2.5 | 2.4 | 37.0 |
| *T. shepardi* | 12.4 | 11.6 | 8.3 | 3.8 | 3.6 | 2.8 | 42.7 |
| *T.californicum* | 16.4 | 13.2 | 8.1 | 4.4 | 2.8 | 3.1 | 48.2 |
| *T. douglasi* | 13.2 | 11 | 8.8 | 4.3 | 2.8 | 2.9 | 43.1 |
| *T. poppensis* | 12.5 | 12.1 | 7.1 | 2.9 | 2.8 | 2.7 | 40.2 |
| *T. monikensis* | 13.8 | 12.6 | 9.6 | 3.4 | 4.2 | 3 | 46.6 |
| *T. cristinae* | 13.7 | 10.9 | 10 | 4 | 3.6 | 3 | 45.3 |
| *T. genevievae* | 15 | 13.4 | 4.3 | 2.5 | 5.4 | 2.8 | 43.5 |
| *T. podura* | 15.7 | 10.8 | 3.1 | 3.1 | 2.6 | 2.3 | 37.7 |

**SM Table 3. Genome assembly statistics**

Genome assembly statistics of sequenced *Timema* species. Haploid genome size represents the estimate from genome profiling of raw reads using Genomescope (*23*). Total sum represents the sum of all scaffolds. The BUSCO score (*22*) is the percentage of conserved single copy orthologs among insects. N is the percentage of unknown nucleotides (gaps) in the assembly. Genes are the number of annotated protein coding genes. Red species reproduce sexually, blue species through parthenogenesis. Although the sequencing coverage was similar across the ten sequenced species (approximately 40x, SM Table 2), all five parthenogenetic species had both higher continuity (NG50 61.9 - 147.4 kbp for parthenogens, vs. 2.1 - 76.1 kbp for sexuals) and higher completeness (97.2 - 98.3% BUSCO genes in parthenogens, vs. 86.4 - 97.2% in sexuals), likely because of systematic differences in heterozygosity between species with different reproductive modes (see main text).

| species | Haploid genome size [Gpb] | Σ [Gpb] | BUSCO [%] | Ns [%] | Genes |
|---|---|---|---|---|---|
| *T. tahoe* | 1.13 | 1.093 | 97.5 | 2.4 | 12771 |
| *T. bartmani* | 1.15 | 1.109 | 97.2 | 2.6 | 14066 |
| *T. shepardi* | 1.23 | 1.153 | 97.2 | 1.7 | 14033 |
| *T.californicum* | 1.3 | 1.220 | 94.4 | 1.8 | 14563 |
| *T. douglasi* | 1.26 | 1.124 | 97.2 | 1.6 | 13824 |
| *T. poppensis* | 1.31 | 1.137 | 93.9 | 1.6 | 15605 |
| *T. monikensis* | 1.12 | 1.099 | 98.3 | 1.7 | 12837 |
| *T. cristinae* | 1.11 | 1.178 | 96.9 | 2.3 | 13882 |
| *T. genevievae* | 1.07 | 1.049 | 97.9 | 1.6 | 12009 |
| *T. podura* | 1.04 | 1.105 | 86.4 | 0.4 | 16529 |

**SM Table 4. Numbers of 1:1 orthologs in different sets of *Timema* species**

See external file “SM_Table_4.tsv”.

**SM Table 5. Origin of the genetic variation among genotypes in parthenogenetic populations**

To distinguish between putative ancestral polymorphisms (shared between sexual and parthenogenetic species) and polymorphisms that appeared in the parthenogenetic lineage after the split from the sexual lineage, we used the SNPs generated for heterozygosity estimates via GATK best practices pipeline (*67*) (see Methods) but with less stringent downstream filtering (min 10x coverage). Homologous SNPs within a species pair were identified with MUMmer v4.0.0beta2 (nucmer and dnadiff with default parameters to keep only unique alignments of genome segments, and custom scripts to discard overlapping alignments), using the genome of the parthenogenetic species as the reference and the one from its sexual relative as the query.

| Species pair | Number of positions analyzed | Variable (within and/or between species) | Same variants in both species | Different variants | Variable only in sexual species | Variable only in partheno-genetic species |
|---|---|---|---|---|---|---|
| *T. bartmani*<br>*T. tahoe* | 852224058 | 8945655 | 26137 | 3683201 | 5052559 | 183758 |
| *T. californicum*<br>*T. shepardi* | 725333178 | 12631427 | 51243 | 4752391 | 7604702 | 223091 |
| *T. cristinae*<br>*T. monikensis* | 816642553 | 19793873 | 87370 | 7078329 | 11677904 | 950270 |
| *T. poppensis*<br>*T. douglasi* | 781906596 | 14109700 | 206188 | 8296511 | 3989068 | 1617933 |
| *T. podura*<br>*T. genevievae* | 636577084 | 27365408 | 325 | 5947476 | 21410167 | 7440 |

**SM Table 6. Number of RNA-seq libraries used for genome annotation**

Species are abbreviated as follows: Tbi = *T. bartmani*, Tce = *T. cristinae*, Tps = *T. poppensis*, Tcm = *T. californicum*, Tpa = *T. podura*, Tte = *T. tahoe*, Tms = *T. monikensis*, Tdi = *T. douglasi*, Tsi = *T. shepardi*, and Tge = *T. genevievae*

| Tissue | Library type | Tbi | Tte | Tce | Tms | Tcm | Tsi | Tpa | Tge | Tps | Tdi |
|---|---|---|---|---|---|---|---|---|---|---|---|
| Whole-Body (Female) | Single-end | 6 | 3 | 6 | 3 | 6 | 3 | 6 | 3 | 6 | 3 |
| Whole-Body (Male) | Single-end | 3 | | 3 | | 3 | | 3 | | 3 | |
| Rep. tract (Female) | Single-end | 3 | 3 | 3 | 3 | 3 | 3 | 3 | 3 | 3 | 3 |
| Rep. tract (Male) | Single-end | 3 | | 3 | | 3 | | 3 | | 3 | |
| Heads (Female) | Single-end | 3 | 3 | 3 | 3 | 3 | 3 | 3 | 3 | 3 | 3 |
| Heads (Male) | Single-end | 3 | | 3 | | 3 | | 3 | | 3 | |
| Legs (Female) | Single-end | 3 | 3 | 3 | 3 | 3 | 3 | 3 | 3 | 3 | 3 |
| Legs (Male) | Single-end | 3 | | 3 | | 3 | | 3 | | 3 | |
| Juvenile (Female) | Paired-end | | | | | 3 | 3 | | | | |
| Juvenile (Male) | Paired-end | | | | | 3 | | | | | |
| Hatchling (Unknown) | Paired-end | | | 7 | 3 | 6 | 3 | 5 | | | 3 |

**SM Table 7A. Accession numbers for raw reads of reference individuals**

Species are abbreviated as follows: Tbi = *T. bartmani*, Tce = *T. cristinae*, Tps = *T. poppensis*, Tcm = *T. californicum*, Tpa = *T. podura*, Tte = *T. tahoe*, Tms = *T. monikensis*, Tdi = *T. douglasi*, Tsi = *T. shepardi*, and Tge = *T. genevievae*

| Library Name | Sp ID | Sample ID | Insert size | SRA sample accession | SRA run accession | Assembly | Genome profiling | Variants |
|---|---|---|---|---|---|---|---|---|
| HYI-7_125 | 4_Tte | Tte_00 | 350 | SRS1972401 | SRR5248900 | * | * | |
| HYI-7_150 | 4_Tte | Tte_00 | 350 | SRS1972401 | SRR5248899 | | * | |
| HYI-17 | 4_Tte | Tte_00 | 550 | SRS1972401 | SRR5248898 | * | * | * |
| HYI-51 | 4_Tte | Tte_00 | 700 | SRS1972401 | SRR5248897 | * | * | |
| HYI-37 | 4_Tte | Tte_00 | 3000 | SRS1972401 | SRR5248896 | * | | |
| HYI-47 | 4_Tte | Tte_00 | 5000 | SRS1972401 | SRR5248895 | * | | |
| HYI-18_125 | 4_Tbi | Tbi_00 | 350 | SRS1972400 | SRR5248892 | * | * | |
| HYI-8_125 | 4_Tbi | Tbi_00 | 350 | SRS1972400 | SRR5248894 | | * | |
| HYI-8_150 | 4_Tbi | Tbi_00 | 350 | SRS1972400 | SRR5248893 | * | * | * |
| HYI-28 | 4_Tbi | Tbi_00 | 700 | SRS1972400 | SRR5248891 | * | * | |
| HYI-38 | 4_Tbi | Tbi_00 | 3000 | SRS1972400 | SRR5248890 | * | | |
| HYI-48 | 4_Tbi | Tbi_00 | 5000 | SRS1972400 | SRR5248889 | * | | |
| HYI-4_125 | 2_Tsi | Tsi_00 | 350 | SRS1972405 | SRR5248924 | * | * | |
| HYI-4_150 | 2_Tsi | Tsi_00 | 350 | SRS1972405 | SRR5248923 | | * | |
| HYI-14 | 2_Tsi | Tsi_00 | 550 | SRS1972405 | SRR5248922 | * | * | * |
| HYI-24 | 2_Tsi | Tsi_00 | 700 | SRS1972405 | SRR5248921 | * | * | |
| HYI-34 | 2_Tsi | Tsi_00 | 3000 | SRS1972405 | SRR5248920 | * | | |
| HYI-44 | 2_Tsi | Tsi_00 | 5000 | SRS1972405 | SRR5248919 | * | | |
| HYI-3_125 | 2_Tcm | Tcm_00 | 350 | SRS1972404 | SRR5248918 | * | * | |
| HYI-3_150 | 2_Tcm | Tcm_00 | 350 | SRS1972404 | SRR5248917 | | * | |
| HYI-13 | 2_Tcm | Tcm_00 | 550 | SRS1972404 | SRR5248916 | * | * | * |
| HYI-23 | 2_Tcm | Tcm_00 | 700 | SRS1972404 | SRR5248915 | * | * | |
| HYI-33 | 2_Tcm | Tcm_00 | 3000 | SRS1972404 | SRR5248914 | * | | |
| HYI-43 | 2_Tcm | Tcm_00 | 5000 | SRS1972404 | SRR5248913 | * | | |
| HYI-5_125 | 3_Tms | Tms_00 | 350 | SRS1972403 | SRR5248912 | * | * | |
| HYI-5_150 | 3_Tms | Tms_00 | 350 | SRS1972403 | SRR5248911 | | * | |
| HYI-15 | 3_Tms | Tms_00 | 550 | SRS1972403 | SRR5248910 | * | * | * |
| HYI-25 | 3_Tms | Tms_00 | 700 | SRS1972403 | SRR5248909 | * | * | |
| HYI-35 | 3_Tms | Tms_00 | 3000 | SRS1972403 | SRR5248908 | * | | |
| HYI-45 | 3_Tms | Tms_00 | 5000 | SRS1972403 | SRR5248907 | * | | |
| HYI-6_125 | 3_Tce | Tce_00 | 350 | SRS1972402 | SRR5248906 | * | * | |
| HYI-6_150 | 3_Tce | Tce_00 | 350 | SRS1972402 | SRR5248905 | | * | |
| HYI-16 | 3_Tce | Tce_00 | 550 | SRS1972402 | SRR5248904 | * | * | * |
| HYI-26 | 3_Tce | Tce_00 | 700 | SRS1972402 | SRR5248903 | * | * | |
| HYI-36 | 3_Tce | Tce_00 | 3000 | SRS1972402 | SRR5248902 | * | | |
| HYI-46 | 3_Tce | Tce_00 | 5000 | SRS1972402 | SRR5248901 | * | | |
| HYI-1_125 | 1_Tdi | Tdi_00 | 350 | SRS1972407 | SRR5248936 | * | * | |
| HYI-1_150 | 1_Tdi | Tdi_00 | 350 | SRS1972407 | SRR5248935 | | * | |
| HYI-11 | 1_Tdi | Tdi_00 | 550 | SRS1972407 | SRR5248934 | * | * | * |
| HYI-21 | 1_Tdi | Tdi_00 | 700 | SRS1972407 | SRR5248933 | * | * | |

| | | | | | | | | |
|---|---|---|---|---|---|---|---|---|
| HYI-31 | 1_Tdi | Tdi_00 | 3000 | SRS1972407 | SRR5248932 | * | | |
| HYI-41 | 1_Tdi | Tdi_00 | 5000 | SRS1972407 | SRR5248931 | * | | |
| HYI-2_125 | 1_Tps | Tps_00 | 350 | SRS1972406 | SRR5248930 | * | * | |
| HYI-2_150 | 1_Tps | Tps_00 | 350 | SRS1972406 | SRR5248929 | | * | |
| HYI-12 | 1_Tps | Tps_00 | 550 | SRS1972406 | SRR5248928 | * | * | * |
| HYI-22 | 1_Tps | Tps_00 | 700 | SRS1972406 | SRR5248927 | * | * | |
| HYI-32 | 1_Tps | Tps_00 | 3000 | SRS1972406 | SRR5248926 | * | | |
| HYI-42 | 1_Tps | Tps_00 | 5000 | SRS1972406 | SRR5248925 | * | | |
| HYI-10_125 | 5_Tge | Tge_00 | 350 | SRS1972399 | SRR5248888 | * | * | |
| HYI-10_150 | 5_Tge | Tge_00 | 350 | SRS1972399 | SRR5248887 | | * | |
| HYI-20 | 5_Tge | Tge_00 | 550 | SRS1972399 | SRR5248886 | * | * | * |
| HYI-53 | 5_Tge | Tge_00 | 700 | SRS1972399 | SRR5248885 | * | * | |
| HYI-40 | 5_Tge | Tge_00 | 3000 | SRS1972399 | SRR5248884 | * | | |
| HYI-50 | 5_Tge | Tge_00 | 5000 | SRS1972399 | SRR5248883 | * | | |
| HYI-9_125 | 5_Tpa | Tpa_00 | 350 | SRS1972398 | SRR5248882 | * | * | |
| HYI-9_150 | 5_Tpa | Tpa_00 | 350 | SRS1972398 | SRR5248881 | | * | |
| HYI-19 | 5_Tpa | Tpa_00 | 550 | SRS1972398 | SRR5248880 | * | * | * |
| HYI-52 | 5_Tpa | Tpa_00 | 700 | SRS1972398 | SRR5248879 | * | * | |
| HYI-39 | 5_Tpa | Tpa_00 | 3000 | SRS1972398 | SRR5248878 | * | | |
| HYI-49 | 5_Tpa | Tpa_00 | 5000 | SRS1972398 | SRR5248877 | * | | |

**SM Table 7B. Accession numbers for raw reads of resequenced individuals**

Species are abbreviated as follows: Tbi = *T. bartmani*, Tce = *T. cristinae*, Tps = *T. poppensis*, Tcm = *T. californicum*, Tpa = *T. podura*, Tte = *T. tahoe*, Tms = *T. monikensis*, Tdi = *T. douglasi*, Tsi = *T. shepardi*, and Tge = *T. genevievae*

| Library Name | Species ID | Sample ID | SRA sample accession | SRA run accession |
|---|---|---|---|---|
| ReSeq_Te07 | 4_Tte | Tte_01 | SRS7638306 | SRR12928425, SRR12928426, SRR12928429-SRR12928438, SRR12928440-SRR12928449 |
| ReSeq_Te08 | 4_Tte | Tte_02 | SRS7638305 | SRR12928399-SRR12928404, SRR12928406-SRR12928415, SRR12928417-SRR12928424 |
| ReSeq_Te09 | 4_Tte | Tte_03 | SRS7638326 | SRR12928367-SRR12928371, SRR12928373-SRR12928382, SRR12928384-SRR12928393, SRR12928395-SRR12928398 |
| ReSeq_Te10 | 4_Tte | Tte_04 | SRS7638327 | SRR12928340-SRR12928349, SRR12928351-SRR12928360, SRR12928362-SRR12928366 |
| ReSeq_Te11 | 4_Tte | Tte_05 | SRS7638328 | SRR12928311-SRR12928315, SRR12928318-SRR12928327, SRR12928329-SRR12928338 |
| CC86B | 4_Tbi | Tbi_01 | SRS7637496 | SRR12928843-SRR12928847, SRR12928849-SRR12928858, SRR12928860 |
| CC86C | 4_Tbi | Tbi_02 | SRS7637495 | SRR12928821-SRR12928824, SRR12928826-SRR12928835, SRR12928838-SRR12928842 |
| CC87B | 4_Tbi | Tbi_03 | SRS7637498 | SRR12928490-SRR12928493, SRR12928495-SRR12928504, SRR12928506-SRR12928515, |

| | | | | |
|---|---|---|---|---|
| | | | | SRR12928517-SRR12928520, SRR12928818, SRR12928820 |
| CC87C | 4_Tbi | Tbi_04 | SRS7638307 | SRR12928468-SRR12928471, SRR12928473-SRR12928482, SRR12928484-SRR12928489 |
| CC88B | 4_Tbi | Tbi_05 | SRS7638309 | SRR12928451-SRR12928460, SRR12928462-SRR12928467 |
| ReSeq_Si01 | 2_Tsi | Tsi_01 | SRS7638289 | SRR12928651-SRR12928659, SRR12928661-SRR12928663 |
| ReSeq_S14 | 2_Tsi | Tsi_02 | SRS7638288 | SRR12928664-SRR12928670, SRR12928672-SRR12928676 |
| ReSeq_Si03 | 2_Tsi | Tsi_03 | SRS7638287 | SRR12928635-SRR12928637, SRR12928639-SRR12928648, SRR12928650 |
| ReSeq_Si16 | 2_Tsi | Tsi_04 | SRS7638284 | SRR12928621-SRR12928626, SRR12928628-SRR12928634 |
| ReSeq_Si18 | 2_Tsi | Tsi_05 | SRS7638285 | SRR12928604, SRR12928606-SRR12928615, SRR12928617-SRR12928620 |
| HM217 | 2_Tcm | Tcm_01 | SRS7638279 | SRR12928757-SRR12928759, SRR12928761-SRR12928770, SRR12928772-SRR12928778 |
| HM218 | 2_Tcm | Tcm_02 | SRS7638277 | SRR12928735-SRR12928737, SRR12928739-SRR12928748, SRR12928750-SRR12928756 |
| HM219 | 2_Tcm | Tcm_03 | SRS7638281 | SRR12928713-SRR12928715, SRR12928717-SRR12928726, SRR12928728-SRR12928734 |
| HM220 | 2_Tcm | Tcm_04 | SRS7638282 | SRR12928695-SRR12928703, SRR12928706-SRR12928712 |

| | | | | |
|---|---|---|---|---|
| HM221 | 2_Tcm | Tcm_05 | SRS7638286 | SRR12928677-SRR12928681, SRR12928683-SRR12928692, SRR12928694 |
| ReSeq_Ms01 | 3_Tms | Tms_01 | SRS7637486 | SRR12928998-SRR12929002, SRR12929004-SRR12929013, SRR12929015 |
| ReSeq_Ms02 | 3_Tms | Tms_02 | SRS7637485 | SRR12928916, SRR12928918-SRR12928924, SRR12928988, SRR12928990, SRR12928991, SRR12928993-SRR12928997 |
| ReSeq_Ms03 | 3_Tms | Tms_03 | SRS7637493 | SRR12928896, SRR12928898-SRR12928905, SRR12928907-SRR12928915 |
| MS_Alp03b | 3_Tms | Tms_04 | SRS7637467 | SRR12929069-SRR12929077, SRR12929080-SRR12929089, SRR12929091, SRR12929092 |
| MS_Alp04b | 3_Tms | Tms_05 | SRS7637463 | SRR12929016, SRR12929048-SRR12929055, SRR12929057-SRR12929066, SRR12929068 |
| CC22B | 3_Tce | Tce_01 | SRS7638290 | SRR12928577-SRR12928581, SRR12928583-SRR12928592, SRR12928595-SRR12928603 |
| CC22C | 3_Tce | Tce_02 | SRS7638291 | SRR12928555-SRR12928559, SRR12928561-SRR12928570, SRR12928572-SRR12928576 |
| CC24B | 3_Tce | Tce_03 | SRS7638292 | SRR12928533-SRR12928537, SRR12928539-SRR12928548, SRR12928550-SRR12928554 |

| | | | | |
|---|---|---|---|---|
| CC24C | 3_Tce | Tce_04 | SRS7637466 | SRR12928521-SRR12928526, SRR12928528-SRR12928532, SRR12928819, SRR12929111, SRR12929113-SRR12929115 |
| CC25B | 3_Tce | Tce_05 | SRS7637461 | SRR12929093-SRR12929100, SRR12929102-SRR12929110 |
| ReSeq_Di02 | 1_Tdi | Tdi_01 | SRS7637469 | SRR12928239, SRR12928250, SRR12928261, SRR12928272, SRR12928283, SRR12928294, SRR12928305, SRR12928961, SRR12928972, SRR12928983, SRR12929022, SRR12929034, SRR12929045 |
| ReSeq_Di04 | 1_Tdi | Tdi_02 | SRS7637489 | SRR12928865-SRR12928870, SRR12928872, SRR12928878, SRR12928889, SRR12928928, SRR12928939, SRR12928950 |
| ReSeq_Di06 | 1_Tdi | Tdi_03 | SRS7637497 | SRR12928806-SRR12928814, SRR12928861-SRR12928864 |
| ReSeq_Di08 | 1_Tdi | Tdi_04 | SRS7638280 | SRR12928792, SRR12928794-SRR12928803, SRR12928805 |
| ReSeq_Di10 | 1_Tdi | Tdi_05 | SRS7638278 | SRR12928779-SRR12928781, SRR12928783-SRR12928791 |
| ReSeq_Ps14 | 1_Tps | Tps_01 | SRS7637462 | SRR12928527, SRR12928538, SRR12928549, SRR12928560, SRR12928571, SRR12928582, SRR12928593, SRR12928605, SRR12929014, SRR12929056, SRR12929067, SRR12929078, SRR12929090, SRR12929101, SRR12929112 |

| | | | | |
|---|---|---|---|---|
| ReSeq_Ps16 | 1_Tps | Tps_02 | SRS7637490 | SRR12928483, SRR12928494, SRR12928505, SRR12928516, SRR12928825, SRR12928836, SRR12928848, SRR12928859, SRR12928906, SRR12928917, SRR12928992, SRR12929003 |
| ReSeq_Ps18 | 1_Tps | Tps_03 | SRS7638308 | SRR12928316, SRR12928328, SRR12928339, SRR12928350, SRR12928361, SRR12928372, SRR12928383, SRR12928394, SRR12928405, SRR12928416, SRR12928427, SRR12928439, SRR12928450, SRR12928461, SRR12928472 |
| ReSeq_Ps08 | 1_Tps | Tps_04 | SRS7637470 | SRR12928317, SRR12928428, SRR12928594, SRR12928705, SRR12928782, SRR12928793, SRR12928804, SRR12928815-SRR12928817, SRR12928837, SRR12928871, SRR12929023, SRR12929079 |
| ReSeq_Ps12 | 1_Tps | Tps_05 | SRS7638283 | SRR12928616, SRR12928627, SRR12928638, SRR12928649, SRR12928660, SRR12928671, SRR12928682, SRR12928693, SRR12928704, SRR12928716, SRR12928727, SRR12928738, SRR12928749, SRR12928760, SRR12928771 |
| CC59_A | 5_Tge | Tge_01 | SRS7637468 | SRR12928980-SRR12928982, SRR12928984-SRR12928987, SRR12928989, SRR12929017-SRR12929021, SRR12929024-SRR12929026 |

| | | | | |
|---|---|---|---|---|
| CC59_C | 5_Tge | Tge_02 | SRS7637484 | SRR12928958-SRR12928960, SRR12928962-SRR12928971, SRR12928973-SRR12928979 |
| CC65_B | 5_Tge | Tge_03 | SRS7637488 | SRR12928937, SRR12928938, SRR12928940-SRR12928949, SRR12928951-SRR12928957 |
| CC66_A | 5_Tge | Tge_04 | SRS7637487 | SRR12928892-SRR12928895, SRR12928897, SRR12928925-SRR12928927, SRR12928929-SRR12928936 |
| CC67_A | 5_Tge | Tge_05 | SRS7637494 | SRR12928873-SRR12928877, SRR12928879-SRR12928888, SRR12928890, SRR12928891 |
| Pa_AB | 5_Tpa | Tpa_01 | SRS7637465 | SRR12929027-SRR12929033, SRR12929035-SRR12929043 |
| PA_CD | 5_Tpa | Tpa_02 | SRS7638329 | SRR12928245-SRR12928249, SRR12928251-SRR12928260, SRR12928262, SRR12928263 |
| PA_E | 5_Tpa | Tpa_03 | SRS7637464 | SRR12928231-SRR12928238, SRR12928240-SRR12928244, SRR12929044, SRR12929046, SRR12929047 |
| H54 | 5_Tpa | Tpa_04 | SRS7638331 | SRR12928293, SRR12928295-SRR12928304, SRR12928306-SRR12928310 |
| H56 | 5_Tpa | Tpa_05 | SRS7638330 | SRR12928264-SRR12928271, SRR12928273-SRR12928282, SRR12928284-SRR12928292 |

**SM Table 8. GO terms enriched in positively selected genes**

Few GO terms are enriched in positively selected genes in each species. This may be partly due to the difficulty in obtaining functional annotations in *Timema*, due to their evolutionary distance from a well characterised insect model system. Species are abbreviated as follows: Tbi = *T. bartmani*, Tce = *T. cristinae*, Tps = *T. poppensis*, Tcm = *T. californicum*, Tpa = *T. podura*, Tte = *T. tahoe*, Tms = *T. monikensis*, Tdi = *T. douglasi*, Tsi = *T. shepardi*, and Tge = *T. genevievae*

| GO ID | Term | Annotated | Significant | Expected | p | sp |
|---|---|---|---|---|---|---|
| GO:0007399 | nervous system development | 315 | 12 | 5.08 | 0.0028 | Tte |
| GO:0006338 | chromatin remodeling | 20 | 3 | 0.32 | 0.0035 | Tte |
| GO:0007476 | imaginal disc-derived wing morphogenesis | 45 | 4 | 0.73 | 0.0055 | Tte |
| GO:0050775 | positive regulation of dendrite morphogenesis | 30 | 3 | 0.48 | 0.0118 | Tte |
| GO:0030178 | negative regulation of Wnt signaling pathway | 31 | 3 | 0.5 | 0.0129 | Tte |
| GO:0043039 | tRNA aminoacylation | 12 | 2 | 0.19 | 0.0152 | Tte |
| GO:0031935 | regulation of chromatin silencing | 14 | 2 | 0.23 | 0.0205 | Tte |
| GO:0008593 | regulation of Notch signaling pathway | 14 | 2 | 0.23 | 0.0205 | Tte |
| GO:0045931 | positive regulation of mitotic cell cycle | 15 | 2 | 0.24 | 0.0234 | Tte |
| GO:0006030 | chitin metabolic process | 17 | 2 | 0.27 | 0.0297 | Tte |
| GO:0009058 | biosynthetic process | 454 | 10 | 7.33 | 0.0306 | Tte |
| GO:0060966 | regulation of gene silencing by RNA | 16 | 2 | 0.26 | 0.0315 | Tte |
| GO:0046331 | lateral inhibition | 44 | 3 | 0.71 | 0.0329 | Tte |
| GO:0007155 | cell adhesion | 47 | 3 | 0.76 | 0.0389 | Tte |
| GO:0007286 | spermatid development | 20 | 2 | 0.32 | 0.0403 | Tte |
| GO:0006997 | nucleus organization | 20 | 2 | 0.32 | 0.0403 | Tte |
| GO:0032990 | cell part morphogenesis | 127 | 7 | 2.05 | 0.0435 | Tte |
| GO:0048814 | regulation of dendrite morphogenesis | 33 | 4 | 0.53 | 0.0451 | Tte |
| GO:0002064 | epithelial cell development | 84 | 4 | 1.36 | 0.0459 | Tte |
| GO:0032259 | methylation | 30 | 3 | 0.33 | 0.0011 | Tbi |
| GO:0007631 | feeding behavior | 12 | 2 | 0.13 | 0.007 | Tms |
| GO:0007450 | dorsal/ventral pattern formation, imaginal disc | 12 | 2 | 0.13 | 0.007 | Tms |
| GO:0016485 | protein processing | 20 | 2 | 0.22 | 0.019 | Tms |
| GO:0007601 | visual perception | 22 | 2 | 0.24 | 0.023 | Tms |
| GO:0007088 | regulation of mitotic nuclear division | 22 | 2 | 0.24 | 0.023 | Tms |
| GO:0031667 | response to nutrient levels | 29 | 3 | 0.31 | 0.03 | Tms |
| GO:0007623 | circadian rhythm | 29 | 2 | 0.31 | 0.039 | Tms |
| GO:0110116 | regulation of compound eye photoreceptor cell differentiation | 31 | 3 | 0.44 | 0.0019 | Tsi |
| GO:0045732 | positive regulation of protein catabolic process | 10 | 2 | 0.14 | 0.0084 | Tsi |
| GO:0043269 | regulation of ion transport | 12 | 2 | 0.17 | 0.0121 | Tsi |
| GO:0031331 | positive regulation of cellular catabolic process | 15 | 2 | 0.21 | 0.0187 | Tsi |
| GO:0045466 | R7 cell differentiation | 17 | 2 | 0.24 | 0.0237 | Tsi |
| GO:0016197 | endosomal transport | 17 | 2 | 0.24 | 0.0237 | Tsi |
| GO:0044773 | mitotic DNA damage checkpoint | 19 | 2 | 0.27 | 0.0279 | Tsi |

| | | | | | | |
|---|---|---|---|---|---|---|
| GO:0035088 | establishment or maintenance of apical/basal cell polarity | 21 | 2 | 0.3 | 0.0354 | Tsi |
| GO:0006520 | cellular amino acid metabolic process | 32 | 2 | 0.46 | 0.0418 | Tsi |
| GO:0006261 | DNA-dependent DNA replication | 13 | 2 | 0.18 | 0.013 | Tcm |
| GO:0098869 | cellular oxidant detoxification | 14 | 2 | 0.2 | 0.016 | Tcm |
| GO:1903008 | organelle disassembly | 14 | 2 | 0.2 | 0.016 | Tcm |
| GO:0007052 | mitotic spindle organization | 14 | 2 | 0.2 | 0.016 | Tcm |
| GO:0003007 | heart morphogenesis | 10 | 2 | 0.2 | 0.016 | Tdi |
| GO:0071985 | multivesicular body sorting pathway | 10 | 2 | 0.2 | 0.016 | Tdi |
| GO:0044262 | cellular carbohydrate metabolic process | 42 | 3 | 0.84 | 0.03 | Tdi |
| GO:0033206 | meiotic cytokinesis | 15 | 2 | 0.3 | 0.039 | Tdi |
| GO:0051046 | regulation of secretion | 21 | 2 | 0.42 | 0.039 | Tdi |
| GO:0072657 | protein localization to membrane | 22 | 2 | 0.44 | 0.039 | Tdi |
| GO:0098656 | anion transmembrane transport | 16 | 2 | 0.32 | 0.039 | Tdi |
| GO:0030001 | metal ion transport | 17 | 2 | 0.34 | 0.044 | Tdi |
| GO:0016197 | endosomal transport | 34 | 3 | 0.68 | 0.048 | Tdi |
| GO:0030855 | epithelial cell differentiation | 81 | 2 | 0.57 | 0.027 | Tge |
| GO:0007030 | Golgi organization | 50 | 2 | 0.35 | 0.046 | Tge |
| GO:0010499 | proteasomal ubiquitin-independent protein catabolic process | 13 | 2 | 0.18 | 0.013 | Tpa |
| GO:0001510 | RNA methylation | 14 | 2 | 0.19 | 0.015 | Tpa |

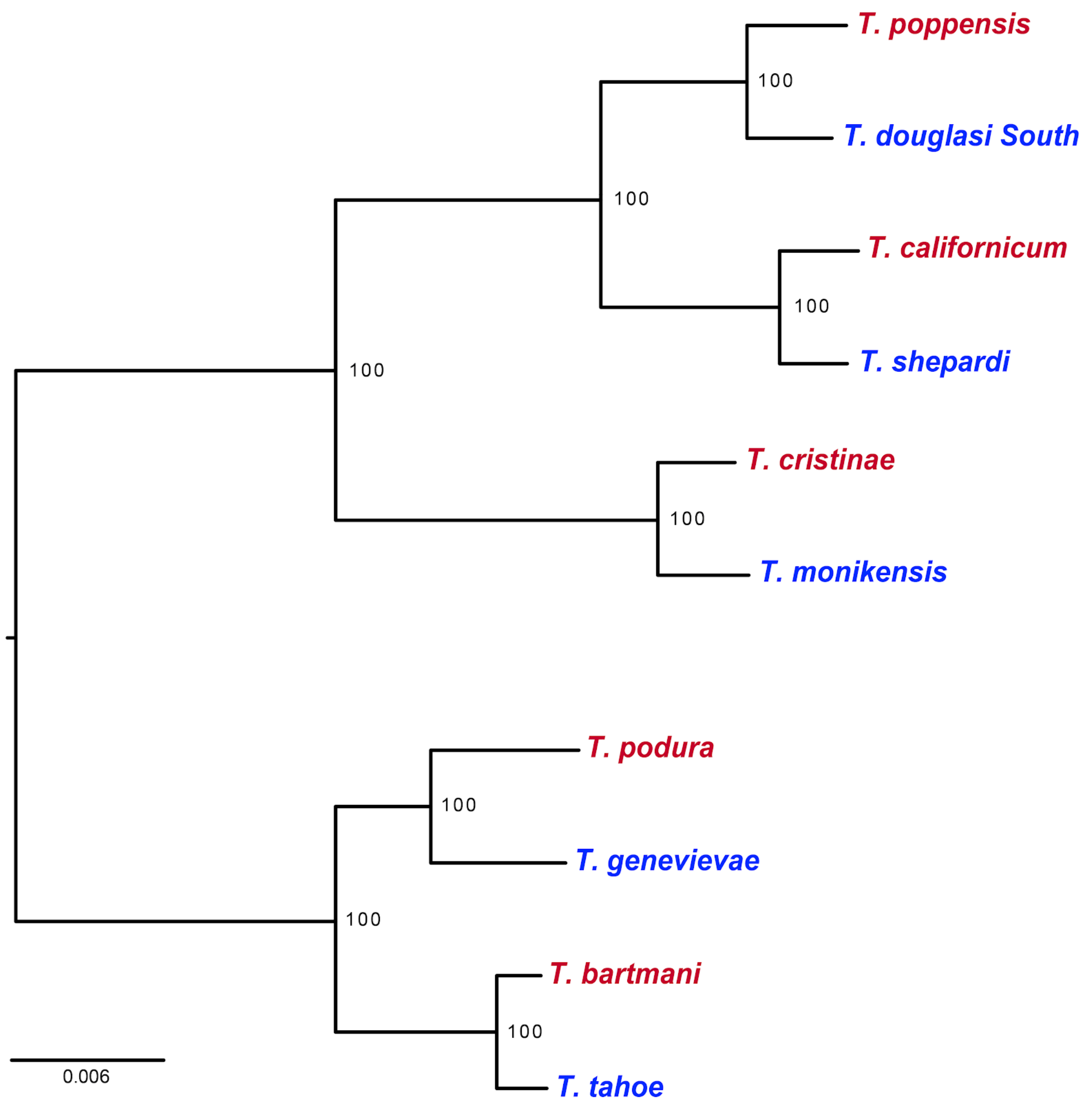


**SM Figure 1** | *Timema* phylogeny. Maximum likelihood tree based on 2377398 orthologous coding DNA positions (from 3975 orthologs), rooted at the midpoint. Branch lengths represent the mean number of substitutions per site. Node labels indicate branch support (%) from 1000 bootstrap replicates. Othologs were aligned using MCoffee (v11.00.8cbe486) (*79*) which was run with the following aligners: mafft_msa, muscle_msa, clustalo_msa (*80*), and t_coffee_msa (*81*). Alignments were concatenated together, and filtered with GBlocks (v. 0.91b, type = codons, minimum block length = 12) to remove large alignment gaps and blocks of Ns (*86*). The tree was then generated with RAxML (*66*), with a GTR+gamma model with 40 rate categories for each codon position.

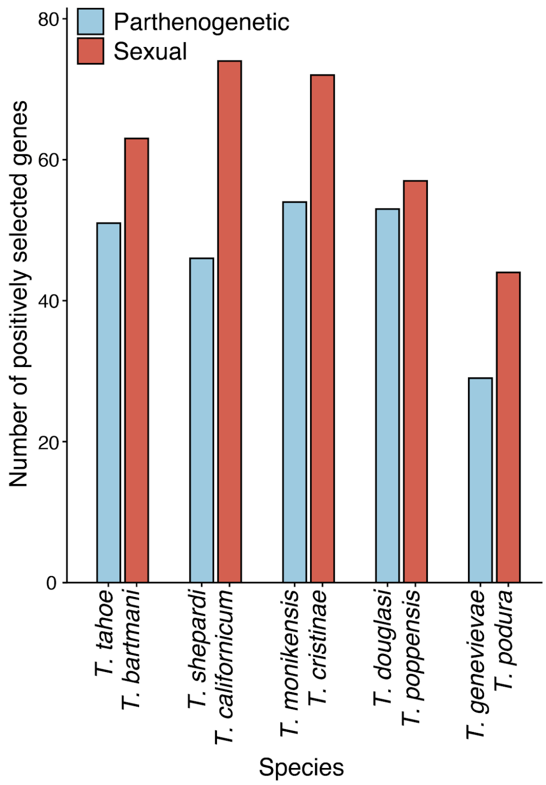


**SM Figure 2** | Number of genes showing evidence for branch-site positive selection on terminal branches with a q-value threshold of 0.01.

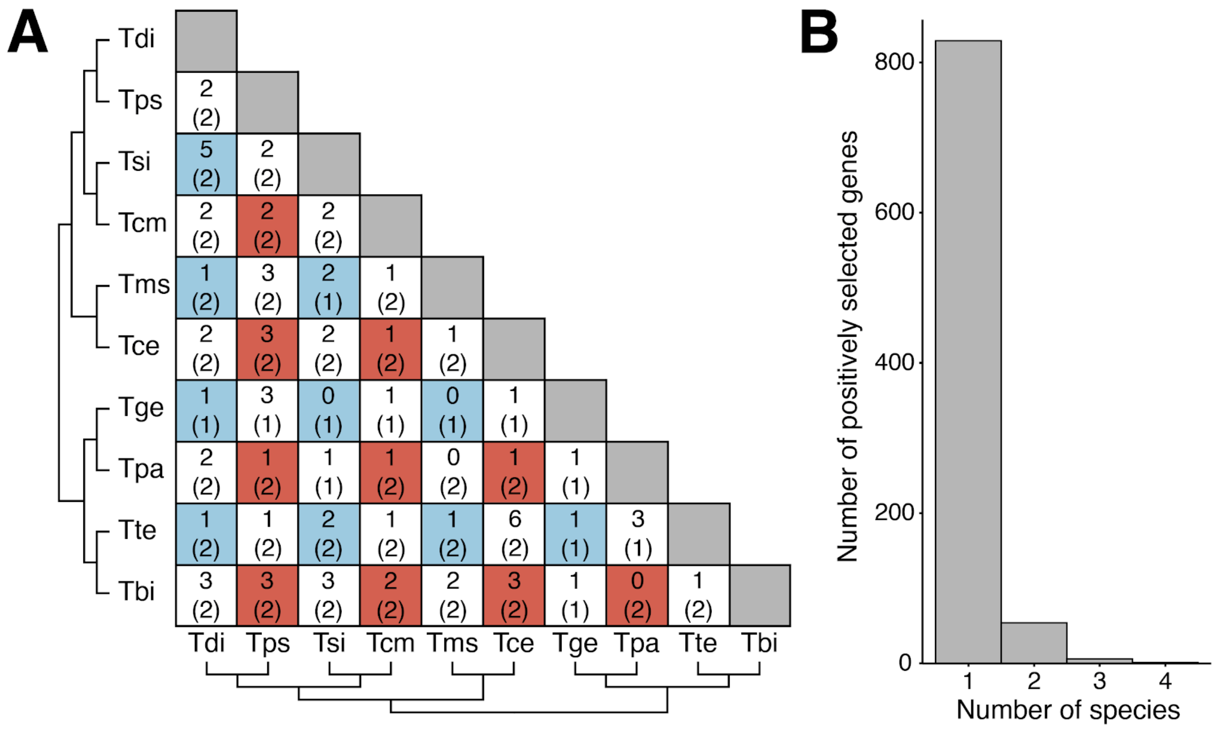


**SM Figure 3** | Positively selected genes are shared between few species. **A.** Matrix showing pairwise overlap of positively selected genes with the number of genes expected by chance given in parentheses. Red cells indicate the overlap between two sexual species, blue between two parthenogenetic species, and white between one sexual and one parthenogenetic species. **B.** Number of species positively selected genes are found in.

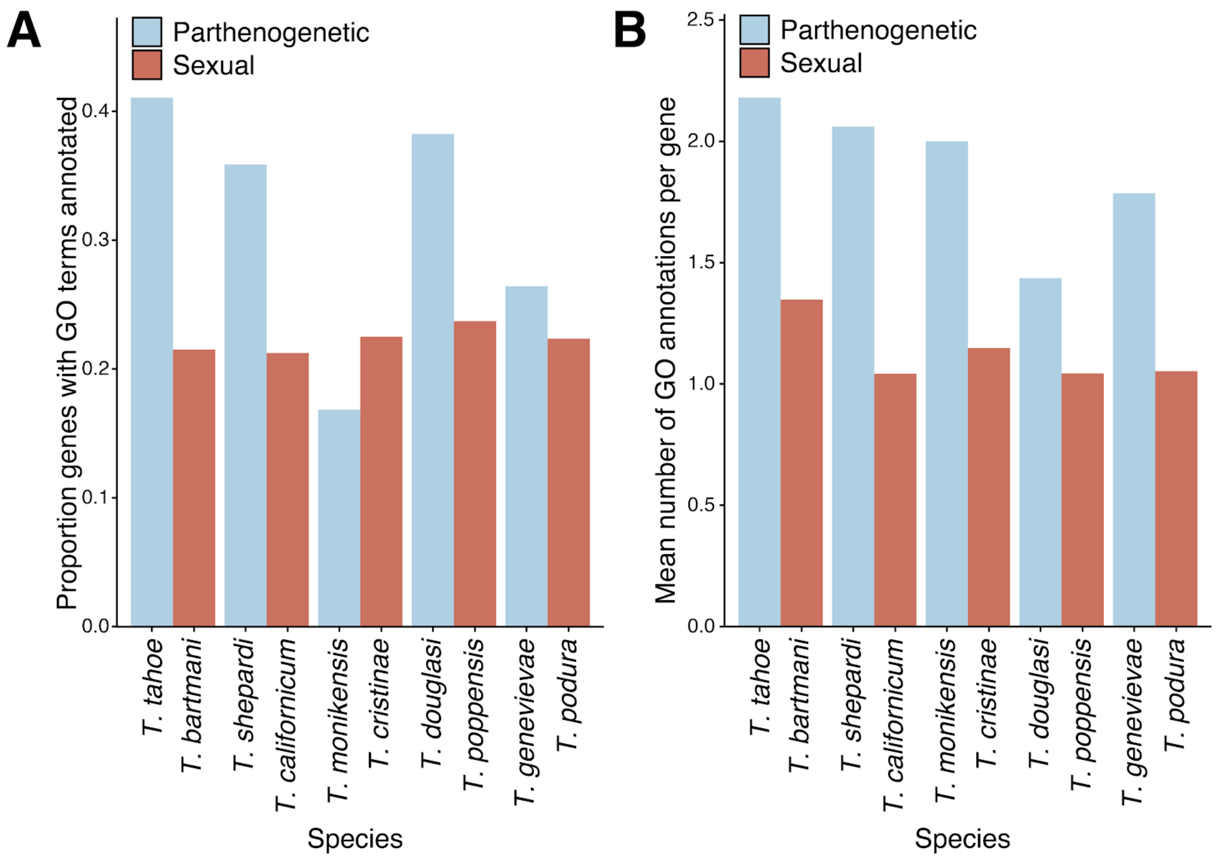


**SM Figure 4** | Positively selected genes in sexual species have fewer annotations than in parthenogenetic species. **A.** Proportion of positively selected genes with at least 1 GO term (biological processes) annotated. **B.** Mean number of GO terms annotated in positively selected genes with at least 1 GO term (biological processes) annotated

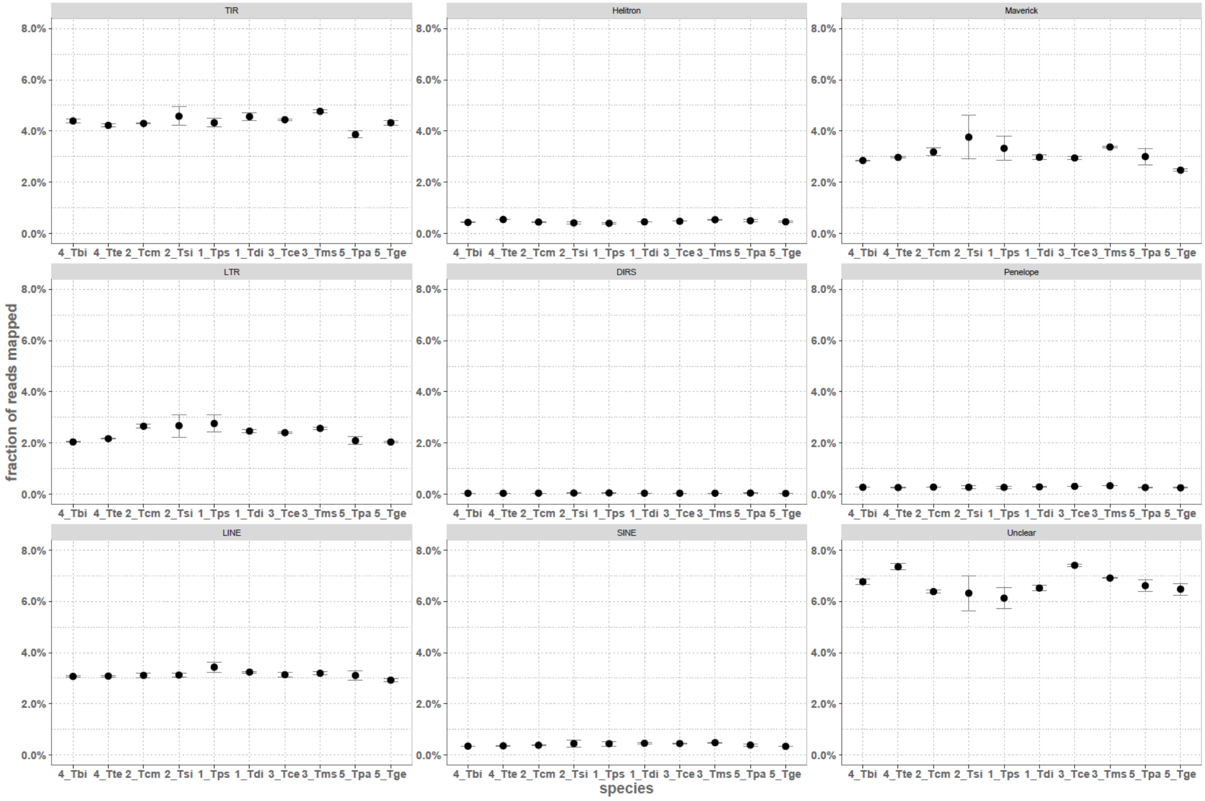


**SM Figure 5 | Genomic transposable element loads, separated by TE orders**.

Error bars represent standard deviation across the six (three for Tsi) sequenced genomes in each species. Species are abbreviated as follows: Tbi = *T. bartmani*, Tce = *T. cristinae*, Tps = *T. poppensis*, Tcm = *T. californicum*, Tpa = *T. podura*, Tte = *T. tahoe*, Tms = *T. monikensis*, Tdi = *T. douglasi*, Tsi = *T. shepardi*, and Tge = *T. genevievae*

**SM text 1: Assembly and annotation pipelines**

Paired-end raw reads were trimmed according to sequencing quality and matched to known Illumina sequencing adapters using Trimmomatic (v.0.36) (*87*). Leading and trailing bases below quality 9 were removed. Reads were scanned using a 4-base sliding window, trimmed when the average quality dropped below 15, and discarded if read length dropped below 96bp (Parameters: PE ILLUMINACLIP: all-adapters.fa:3:25:6 LEADING:9 TRAILING:9 SLIDINGWINDOW:4:15 MINLEN:96). The raw mate-pair reads were de-linked and reverse complemented using NxTrim (v. 0.4.1) (*88*) with the parameter "--preserve-mp". Unlinked pairs without identified adapter sequence, called unknown pairs, were also considered as valid mate pairs as they had a similar distribution of insert sizes as mate pairs with identified linker sequence.

Filtered paired-end reads were *de novo* assembled using ABySS (v. 1.9.0) (*52*, *89*) with default parameters and k-mer sizes predicted to be optimal using kmergenie (*90*). The k-mer sizes were 83, 87, 83, 87, 83, 89, 81, 81, 65 and 87 for *Timema poppensis*, *T. douglasi*, *T. californicum*, *T. shepardi*, *T. cristinae*, *T. monikensis*, *T. barmani, T. tahoe, T. podura and T. genevieve* respectively. Assembled contigs longer than 250 bases were scaffolded using BESST (v. 2.2.5) (*53*) with default parameters and gap-filled with GapCloser (v. 1.12-r6), a module of the SOAP package (*91*).

Genome assemblies were decontaminated using BlobTools (v. 0.9.19.5) (*54*). Hit files were generated after a BlastN (v. 2.6.0) (*74*) against the NBCI nt database (v 2016-06) (*92*), searching for hits with sequence identity above 85% and an e-value below 1e-25 (Parameters: -task megablast -culling_limit 5 -evalue 1e-25 -perc_identity 85). Scaffolds without hits to metazoans were removed from the assemblies. The genome assembly completeness was assessed with BUSCO (v. 3.0.2) (*22*) against the insecta_odb9 lineage and the -long option. For genome annotation, we took a total of 231 publically available RNA-seq libraries for *Timema* from different tissues, sexes and developmental stages as expression evidence (min per species = 12, see SM Table 6) (*37*, *55*, *56*). Before mapping reads to the genomes, adapter sequences were trimmed from raw reads with CutAdapt (v. 1.15) (*93*). Reads were then quality trimmed using Trimmomatic (v. 0.36) (*87*), clipping leading or trailing bases with a phred score of <10 from the read, before using a sliding window from the 5' end to clip the read if 4 consecutive bases had an average phred score of <20. Any reads with a sequence length of <80 after trimming were discarded. All trimmed RNA-seq reads were then mapped against the genomes as single end reads using STAR (v. 2.5.3a) (*94*) under the "2-pass mapping" mode and default parameters. The STAR outputs were then used to produce transcriptome assemblies using Trinity (v. 2.4.0) (*57*) "genome guided" mode (Parameters: --genome_guided_max_intron 100000 --SS_lib_type R). Finally, the transcriptome assemblies were filtered following Trinity developers recommendations (https://github.com/trinityrnaseq/trinityrnaseq/wiki/Trinity-FAQ): Briefly, filtered RNA-seq reads were mapped back against the transcriptomes using Kallisto (v. 0.43.0) (*95*) with options --bias and --single, then transcripts with at least 1 TPM in any sample were retained.

Genome scaffolds >1000 bp were annotated, protein coding genes were predicted using MAKER (v. 2.31.8) (*58*) in a 2-step iterative way as described in Campbell *et al.* (*96*) with minor modifications following author recommendations. For the first iteration, genes were predicted using Augustus (v. 3.2.3) (*97*) trained with the BUSCO results. A combination of UniProtKB/Swiss-Prot (release 2018_01) (*98*) and the BUSCO insecta_odb9 proteome were used as protein evidence. The Trinity assembled RNA-seq reads (described above) were used as transcript evidence. The resulting gene models were then used to retrain Augustus as well as SNAP (v. 2013.11.29) (*99*) and a second iteration was performed. Predicted protein coding genes were then functionally annotated using Blast2GO v5.5.1 (*100*, *101*) with default parameters against both the NCBI non-redundant arthropods protein database, and the *Drosophila melanogaster* (drosoph) database, to produce two sets of functional annotations, one derived from all arthropods and one specifically from *Drosophila melanogaster*.

#### SM text 2: Horizontal Gene Transfers (HGTs) are not facilitated by parthenogenesis

Genomic analyses of bdelloid rotifers, a group that likely persisted and diversified in the absence of canonical sex for over 40 million years (*102*), revealed that bdelloids carry an unusually large amount (6.2% - 9.1%) of horizontally acquired genes compared to sexual lophotrochozoans (0.08% - 0.7%) (*17*, *103*–*105*). Unusually high proportions of HGT-derived genes were also identified in parthenogenetic root-knot nematodes (*106*, *107*) and springtails (*108*). These findings led to the suggestion that parthenogenesis might favor the retention of horizontally acquired genes, and may perhaps confer adaptive benefits that could compensate for the absence of recombination and outcrossing (*106*), although such patterns are not shared by most other parthenogenetic animal genomes (*17*). Analyzing HGT events in *Timema* provided no evidence for parthenogenesis facilitating the retention of HGTs. We identified 55 putative HGT events in the 10 *Timema* species, with up to 50 sequences each, for a total of 704 HGT-derived sequences (351 in the five sexual species vs. 353 in the five parthenogenetic species). The genome of each *Timema* species included approximately 70 HGT-derived sequences, comparable to values from metazoa in general (*109*). All putative HGT families were shared by at least six *Timema* species, and only one putative HGT event occurred in a specific clade (HGT family shared between two sexual and two parthenogenetic species of the Northern clade) while all other HGT events were shared between at least two clades of *Timema*.

Of note, out of the 55 HGT families, 34 featured significant similarities with sequences from two plant pathogens (*Phytophthora infestans* and *Pythium ultimum*), and displayed a high-glycine content, due to many ‘GGG’ repeats. This repeated motif is very similar to the loricrin-like protein described in *Phytophthora infestans* by Guo et al. (*110*), which is suggested to be involved in plant infection.

Out of the 21 remaining families, only 4 showed a phylogenetic pattern consistent with an old HGT event, sometimes shared with *Zootermopsis nevadensis* (the closest species in our reference database). However, the terminal branches leading to the putatively-transferred sequences were too long for a reliable identification of the donor species. The phylogenetic evaluation of the other families was not conclusive, as commonly observed in HGT detection studies (e.g. see (*111*)).

**SM text 3: Analysis of heterozygosity for SNPs and SVs**

We present two estimates of heterozygosity, one based on a reference-free technique (kmer spectra analysis using Genomescope (v. 2) (*23*), the other using sequencing reads mapped to reference genomes to call SNPs with GATK ((*67*), see Methods).

The kmer spectra of all sexual species displayed distinct haploid coverage peaks representing heterozygous kmers (SM Figure 6A), contrasting with the kmer spectra of parthenogenetic species, where no distinct peaks were visible (SM Figure 6B). To confirm that no heterozygous kmers were present at the expected haploid coverage in parthenogens, we used Smudgeplot (*23*), a technique to extract closely related kmer pairs representing heterozygous and paralogous kmer pairs. While in the sexual species, kmers from the 1n peak paired together in heterozygous kmer pairs (AB smudge on SM Figure 6C), no diploid kmer pairs were detected in the parthenogenetic species (SM Figure 6D). We conclude that heterozygosity estimates for the parthenogenetic species cannot be based on k-mer spectra analyses because the heterozygosity levels are too low to reliably fit the distribution estimating haploid kmers in the kmer spectra. Unreliable heterozygosity estimates based on k-mer spectra analyses for species with very low heterozygosity was already reported in Jaron et al. (*17*), suggesting that with the current quality of sequencing data, kmer methods do not have resolution for very small heterozygosity levels.

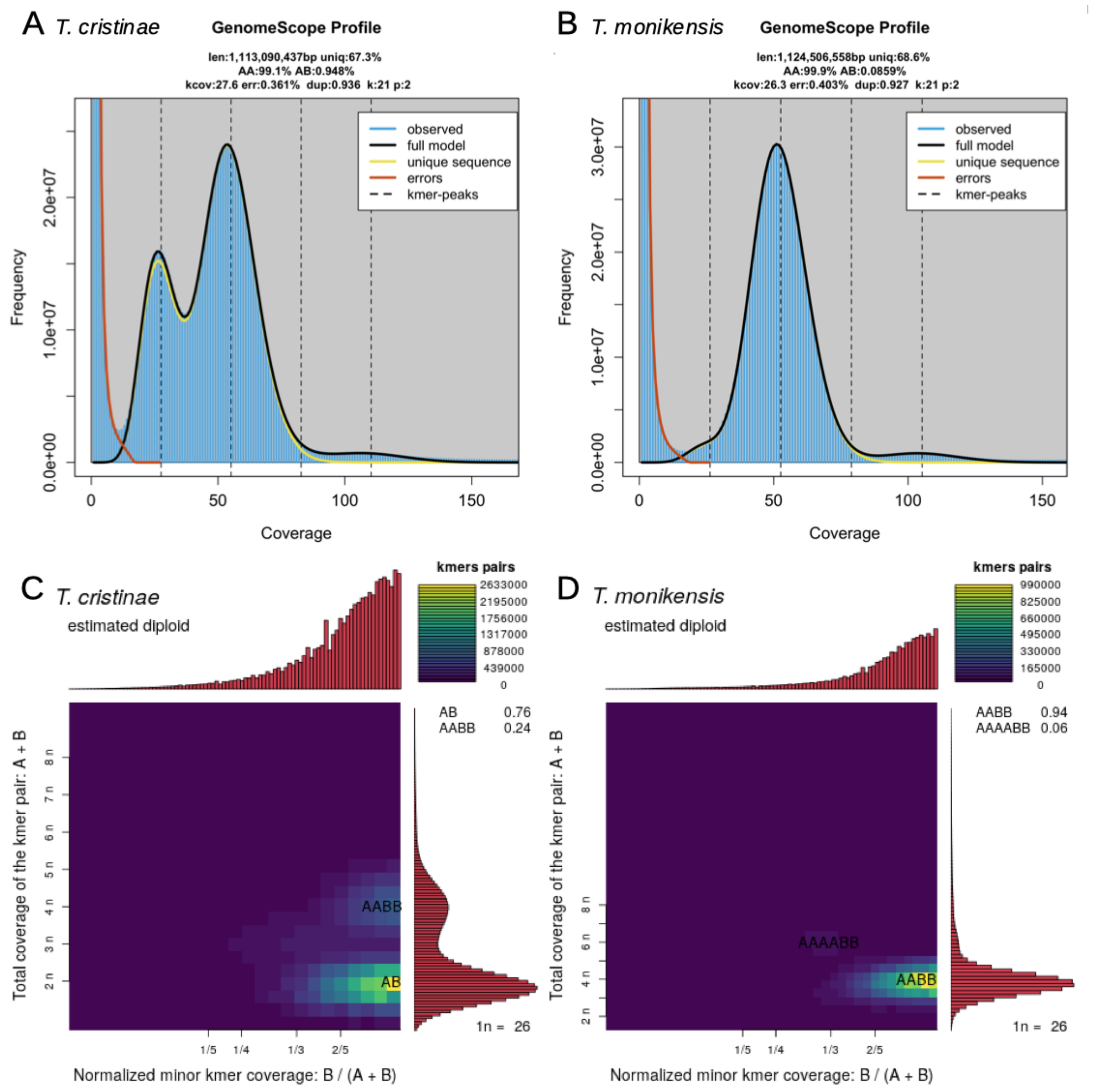


**SM Figure 6**: Genome profiling examples for a sexual (*T. cristinae*, panels A and C) and a parthenogenetic (*T. monikensis*, panels B and D) *Timema* species.

Because we could not estimate heterozygosity of parthenogens using kmer-spectra analyses, we estimated nucleotide heterozygosity using SNP calling. It is important to note, however, that this method generates an underestimation of heterozygosity given our fragmented reference genomes (SM Table 3) and relatively modest coverage (~14 - 21x) of re-sequenced samples. Therefore, our SNP heterozygosity estimates in *Timema* are useful for comparing sexual and parthenogenetic species, but are not accurate estimates of heterozygosity in *Timema* (which range from 0.36

to 2.16% for sexual species, i.e., 2-6 times higher than the SNP-based estimates, Figure 2). In agreement with genome profiling, we find very low, nearly negligible levels of heterozygosity in parthenogenetic species (Figure 2). Furthermore, a large portion of the heterozygous SNP calls in parthenogens  showed an unexpectedly high coverage (SM Figure 7). This excess coverage of heterozygous positions in parthenogens suggests that heterozygous sites in parthenogens largely stem from merged paralogs, further supporting that a very large proportion (or maybe even all) of the called heterozygous variants in parthenogens are just artifacts of the SNP calling pipeline using whole genome data.

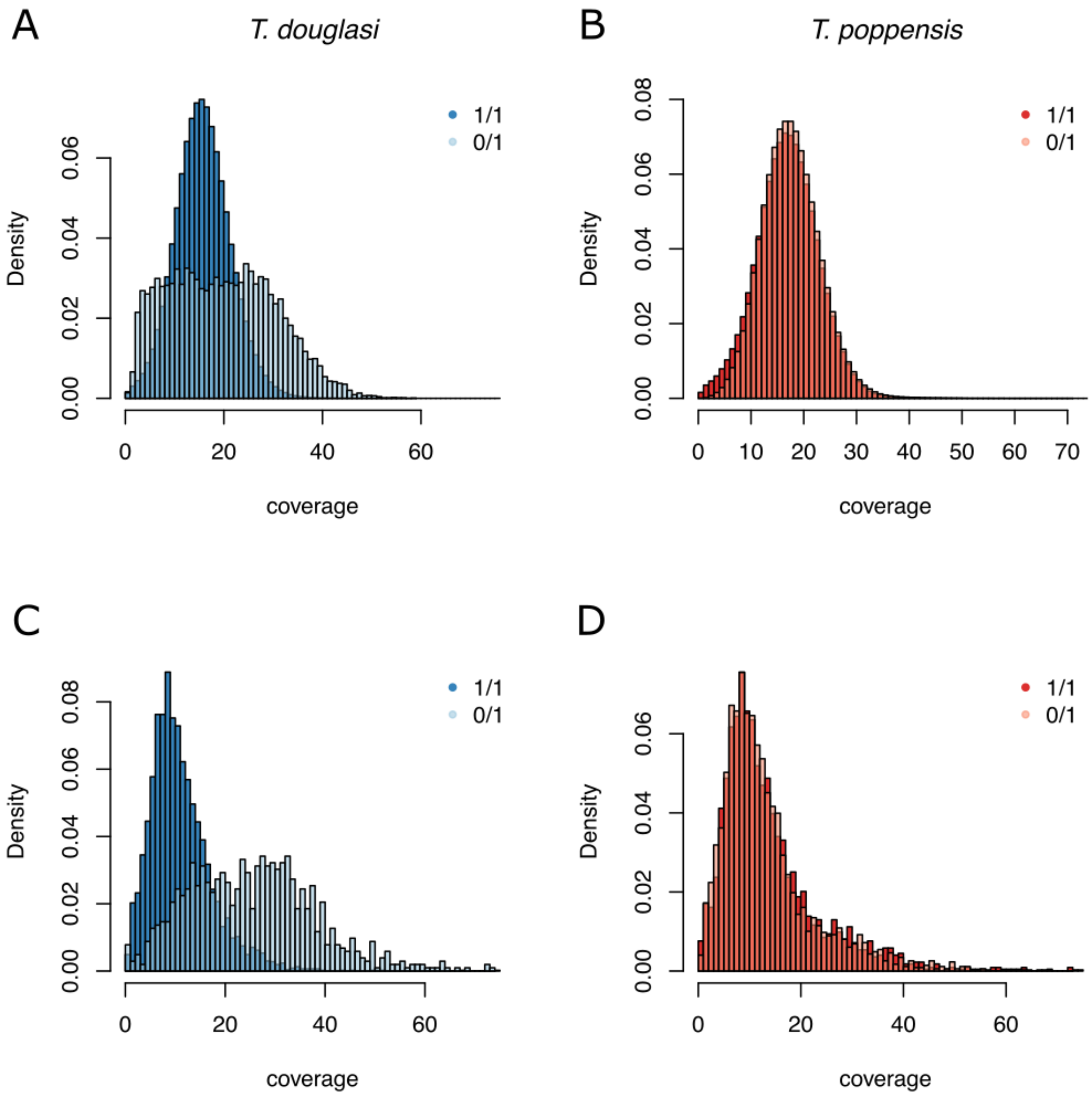


**SM Figure 7**: Densities of coverages supporting **A** SNPs found in the homozygous state (1/1), or heterozygous (0/1) in *T. douglasi*. **B** In sexual *T. poppensis* **C** Densities of split read coverage support of SVs in homozygous or heterozygous states in *T. douglasi* **D**. *T. poppensis*. Both heterozygous SNPs and heterozygous SVs show unexpected coverage distributions in parthenogenetic *T. douglasi* (blue), while coverages supporting SNPs in sexual *T. poppensis* (red) are independent of the genotype. There is a small difference in homozygous and heterozygous SV coverages in sexuals, suggesting that at least some fraction of those heterozygous SVs are also false positives. However, overlap of the two distributions is much greater than in the case of parthenogenetic *T. douglasi* (panel C).

We further investigated if there were any heterozygous structural variations in parthenogenetic *Timema*, as those could be potentially hidden to SNP analysis. Consistent with the previous two analyses, the SV heterozygosity levels were substantially lower in the parthenogens than in their sexual sister species (Figure 2). However, we also detected a non-negligible amount of heterozygous structural variants. We therefore manually curated all heterozygous structural variants found in *T. monikensis* using samplot (v1.0.1) (*112*), but did not find a single variant clearly supported by reads (results not shown). Since structural variant calling from short read data has a high rate of false positives regardless of the method used (*113*), we decided to verify variants using a PacBio long-read dataset (~32x coverage) for one of the parthenogenetic species (*T. douglasi;* PRJNA673001). We assembled the long-read data of *T. douglasi* using Redbean (formerly wtdbg; v2.5) assembler (*114*) with parameters recommended for moderately sized genomes: -L 1000 -x preset3 -g 1300m. This genome assembly was used for SV calling using ngmlr (v0.2.7) and the Sniffles (v1.0.11) pipeline (*115*) with default parameters for SV calling using long read data. In total, we found only 6 heterozygous SVs: 4 deletions and 2 insertions. We visualized the SVs alongside their read support using samplot and found that none were well supported (SM Figure 8) suggesting that the heterozygous SVs called using short read data represent noise in the absence of a signal from real heterozygous SVs.

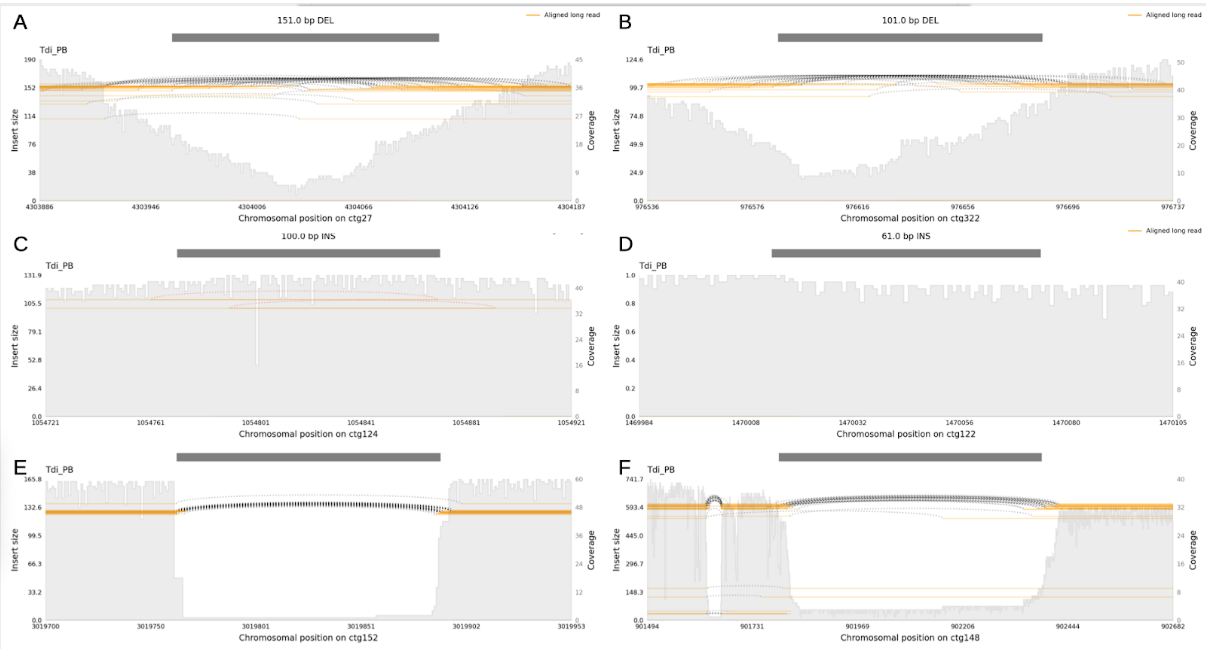


**SM Figure 8.** All 6 heterozygous SVs called in the *T. douglasi* long read dataset. SVs on panels A and B are located in repetitive regions which is causing the uneven distribution of coverages and variable lengths of gaps. Variants on panels C - F are not supported by approximately half of the reads. Variant C is probably due to rare chimeric reads, and variant D does not seem to have any support at all. Conversely, SVs on panels E and F have very low support for the reference sequence. See examples provided in the manual of samplot for comparison to a well supported heterozygous SV.

In conclusion, we used four complementary approaches based on three different data sources: kmer spectra analysis on raw sequencing reads of the reference individuals, SNP and SV heterozygosity estimates using variant calling based on resequencing data, and finally a long read dataset of *T. douglasi*, which was independently assembled and is therefore free of any potential biases introduced in a short read assembly. Our analyses comprehensively show the absence of heterozygous loci in the parthenogenetic *Timema* genome assemblies. Residual heterozygosity could be potentially found in repetitive regions, such as centromeres and telomeres (see also SM text 5), as all our effort to detect heterozygosity focused on alleles with 1n coverage (half of the genome coverage). However, detecting heterozygosity in such regions requires chromosome-scale assemblies based on long-read sequencing technologies, which are currently not available for parthenogenetic *Timema*.

**SM text 4: Locating microsatellite markers in the genome assemblies**

Previous research, based on microsatellite markers, suggested that oogenesis in parthenogenetic *Timema* was functionally mitotic, as there was no loss of heterozygosity between females and their offspring (*18*). Yet our genome data reveal complete or almost complete homozygosity in the genome assemblies of parthenogens (see main text). The most likely reconciliation of these contrasting results is that heterozygosity is maintained in only a small portion of the genome, for example the centromeres or telomeres, or between paralogs.

To investigate these possibilities, we searched for the primer pairs used to amplify the nine microsatellites in the genome assembly v1.3 of the sexual species *T. cristinae* from Nosil et al (*35*). This assembly is currently the most complete and least fragmented *Timema* assembly available, and the microsatellites used by (*18*) were originally developed for *T. cristinae*. We used Blast to find primer pairs <500 bp apart, on opposite strands, and retained significant hits with at least 80% of the primer sequences covered. We then verified whether the retained hits comprised the expected microsatellite repeat motif.

Using this approach, we were able to locate six of the nine microsatellites in the v1.3 assembly (SM Table 9). Two of the six microsatellites had multiple hits in the genome (SM Table 9). In combination, these results support the idea that microsatellite heterozygosity detected in *Timema* parthenogens may be a combination of heterozygosity in centromere or telomere regions (microsatellites not detected in the assembly) and heterozygosity between paralogs (microsatellites with multiple copies in the *T. cristinae* assembly).

**SM Table 9 |** Microsatellites located in the v1.3 genome of *T. cristinae*. Msat name: Microsatellite name from (*18*). Indicated are the scaffolds where a given microsatellite was found (Scaffold), the location of the microsatellite midpoint on the scaffold (Position), the linkage group (LG), the size of the microsatellite in the v1.3 assembly and the expected size range given microsatellite genotypes in *T. cristinae* (Length (expected)), and whether the expected microsatellite repeat motif was present.

| Msat name | Scaffold | LG | Position | Length (expected) [bp] | Motif |
|---|---|---|---|---|---|
| **tim-3 (a)** | CM009483.1 | LG8 | 32962807 | 159 (82-109) | Yes |
| **tim-3 (b)** | CM009476.1 | LG13 | 19984463 | 169 (82-109) | Yes |
| **tim-4** | CM009477.1 | LG2 | 46600506 | 119 (83-125) | Yes |
| **tim-5** | CM009481.1 | LG6 | 7421031 | 198 (124-238) | Yes |
| **tim-6** | CM009474.1 | LG11 | 12218418 | 264 (253-283) | Yes |
| **tim-7** | CM009482.1 | LG7 | 19035853 | 156 (120-195) | Yes |
| **tim-8 (a)** | CM009484.1 | LG9 | 23487739 | 138 (133-148) | Yes |
| **tim-8 (b)** | CM009479.1 | LG4 | 85533694 | 190 (133-148) | No |
| **tim-8 (c)** | CM009478.1 | LG3 | 981498 | 498 (133-148) | No |

#### SM text 5: Polymorphism in parthenogenetic and sexual *Timema* populations

To compare the distribution of polymorphism along different genomes, we mapped population-level variation for SNPs and SVs inferred from 2 to 5 re-sequenced individuals per population to our species-specific reference genomes (see main text). We then anchored our reference genome scaffolds to the 12 autosomal linkage groups of a previously published assembly of the sexual species *T. cristinae* (v1.3 from Nosil et al. (*35*)) using MUMmer (v. 4.0.0beta2) (*70*) (see Methods for details). Note that we excluded LG13, classified as the X chromosome in Nosil et al (*35*), because our detailed analyses of X-chromosomes in *Timema* revealed that LG13 did not correspond to the X (Parker et al in prep). We also removed X-linked scaffolds assigned to autosomal LGs in the v1.3 *T. cristinae* assembly and used this “cleaned” set of linkage groups (referred to as v1.4) in all our analyses with positional information. Depending on the species, we were able to anchor between 59 and 558 Mbp of our genomes to the *T. cristinae* LGs.

We also examined how genetic variation was distributed between individuals by producing phylogenetic trees for the re-sequenced and reference individuals of each sexual-parthenogenetic sister species pair. Sequences for re-sequenced individuals were obtained by mapping reads of each re-sequenced individual to the reference genomes with BWA-MEM (v0.7.15) (*76*). Multi-mapping and poor quality alignments were filtered (removing reads with XA:Z or SA:Z tags or a mapq < 30). We removed PCR duplicates with Picard (v. 2.9.0) (http://broadinstitute.github.io/picard/) and performed indel realignment with GATK (v. 3.7) (*67*). Genomic sequences for each re-sequenced individual were then generated using AngsD (v. 0.921) (-doCounts 1 -doFasta 2) (*116*) with a minimum depth of 5 and a maximum depth of twice the median genome coverage. Coding sequences of 1-to-1 orthologs (2198) were extracted using gffread from the Cufflinks (v. 2.2.1) package (*117*). These sequences were codon-aligned using PRANK (v.100802) (*118*) concatenated together, and filtered with GBlocks (v. 0.91b, type = codons, minimum block length = 12) to remove large alignment gaps and blocks of Ns (*86*). Trees were generated with RAxML (*66*), with a GTR+gamma model with 40 rate categories for each codon position (i.e. each codon position (1st ,2nd, 3rd) was partitioned to allow a distinct model to be fitted to it) to produce an ML tree with 1000 bootstraps.

#### SM text 6: Polymorphism and color morphs on LG8 in the species *T. californicum* and *T. monikensis*

We found very high population polymorphism for both SNPs and SVs on LG8 in *T. californicum* and *T. monikensis* (Figure 3B). A site frequency spectrum (SFS) in *T. californicum* based on the SNPs called in the 5 resequenced individuals revealed that the polymorphism in this species was likely generated by the presence of two distinct haplotypes. The genotype structure for the 5 individuals was similar for a large portion of SNPs on LG8 (1/1 0/0 0/1 0/1 0/1; SM Figure 9A), with three individuals heterozygous, and the two remaining ones homozygous for alternative alleles (SM Figure 9A). The LG8 SNP genotypes further matched the grey versus green color morphs, with the grey morph known to have recessive inheritance (*119*): the individual used to build the reference genome (SNP genotypes 0/0) was grey, as was resequenced individual 2 (Tcm_02) with 0/0 genotypes. The four remaining resequenced individuals (with 1/1 or 0/1 genotypes) were green. The size of the putative haplotypes associated with green or grey morphs is considerable, spanning approximately 24 Mbp on LG8. The genotypes at LG8 in *T. monikensis* were also correlated with color morphs. Brown or beige melanistic morphs featured 0/0 genotypes in the high polymorphism region of LG8, while green individuals had 1/1 genotypes (SM Figure 9B).

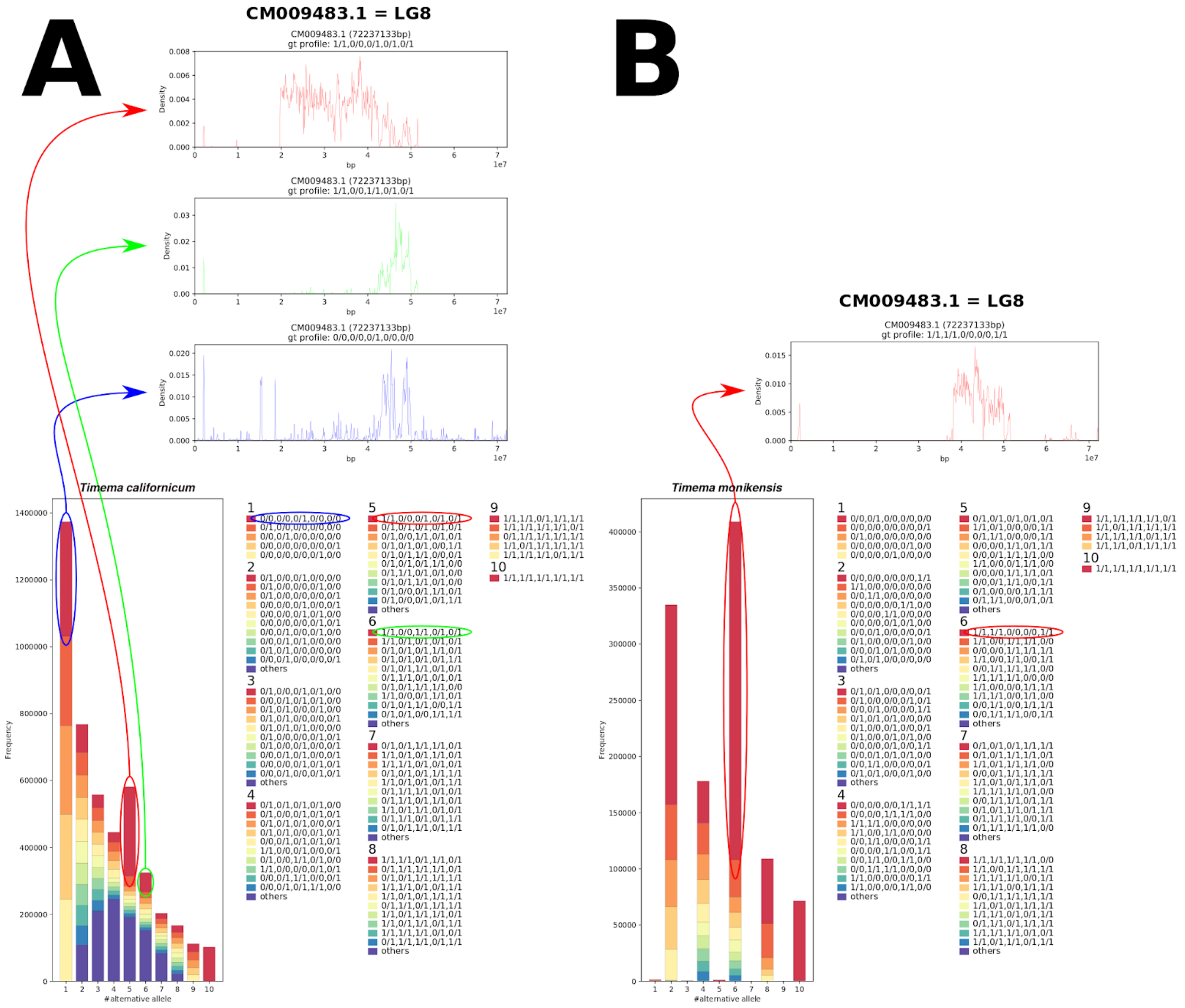

**SM Figure 9.** Site Frequency Spectrum for the five re-sequenced individuals of *T. californicum* (A) and *Timema monikensis* (B), generated with Pop-Con (https://github.com/YoannAnselmetti/Pop-Con), indicating the genotype distributions for each count of alternative alleles. For *T. californicum*, the peak at count 5 is generated by the overrepresented genotype structure 1/1 0/0 0/1 0/1 0/1, and almost all SNPs with this structure (97.34%) map to LG8, suggesting the presence of two divergent haplotypes on LG8. For *T. monikensis*, we observed a similar overrepresentation at allele count 6, for the genotype structure 1/1 1/1 0/0 0/0 1/1, and most of the SNPs with this structure (78.43%) map to LG8 .

We also investigated the presence of an approximately 0.5 Mb deletion on LG8, suggested by Villoutreix et al. (*120*) to determine the green morphs in *T. californicum,* in our ~24 Mbp long haplotypes. Because our reference genome was based on a grey individual (which would be homozygous for the deletion-free haplotype), we expected to observe normal coverage for this region in the re-sequenced grey individual, zero coverage in the green re-sequenced individual homozygous for the alternative haplotype, and half the coverage in the three heterozygous individuals. We observed a coverage reduction in all individuals at the focal region, which could be due to an enrichment in repetitive sequences (non-uniquely mapping reads are not included for coverage estimations). Nevertheless, the grey individual featured somewhat higher coverage, consistent with the deletion suggested by Villoutreix et al. (*120*) (SM Figure 10). Further studies are required to characterize the contribution of the two divergent, ~24 Mbp long haplotypes, and the putative 0.5 Mbp deletion in one of the haplotypes, to color polymorphism in *T. californicum*.

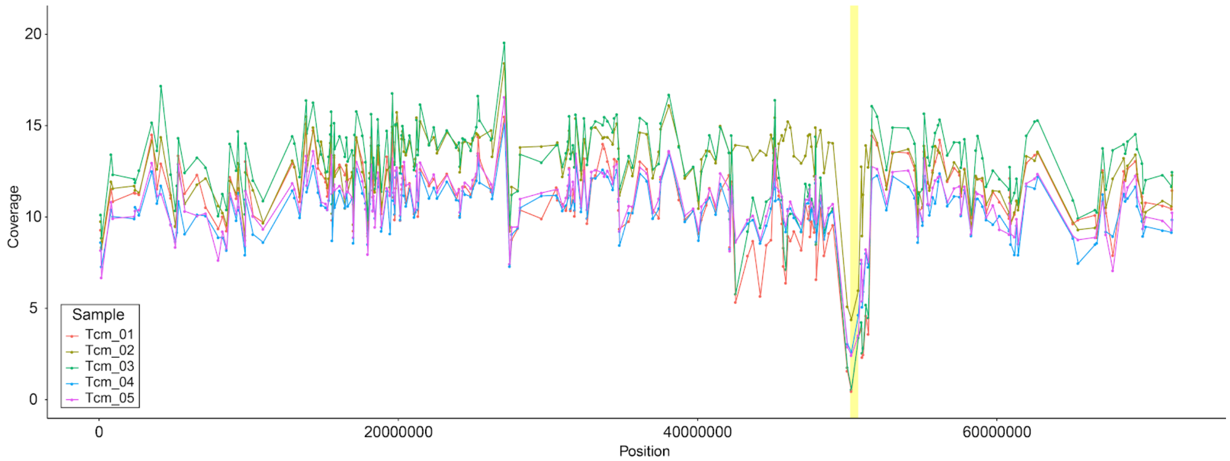


**SM Figure 10.** Coverage along LG8 for 5 resequenced *T. californicum*. Coverage was estimated by mapping reads to the *T. californicum* genome scaffolds, and scaffolds were anchored on *T. cristinae* linkage groups (see methods and SM text 5). The region with the expected deletion is highlighted in yellow. If there was a deletion on LG8 determining the green morph, grey individual Tcm_02 should feature higher coverage than the other individuals which are green.
